# Cell and nuclear size are associated with chromosomal instability and tumorigenicity in cancer cells that undergo whole genome doubling

**DOI:** 10.1101/2025.03.28.645986

**Authors:** Mathew Bloomfield, Sydney Huth, Daniella McCausland, Ron Saad, Nazia Bano, Tran Chau, Megan Sweet, Nicolaas Baudoin, Andrew McCaffrey, Kai Fluet, Eva M. Schmelz, Uri Ben-David, Daniela Cimini

## Abstract

Whole genome doubling (WGD) is a frequent event in cancer evolution associated with chromosomal instability, metastasis, and poor prognosis. While the genomic consequences of WGD are well documented, the effects of non-genetic alterations that accompany WGD, such as changes to cell and nuclear size, on tetraploid (4N) cancer cell physiology are less understood. Here, we show that cell and nuclear volume do not always scale with DNA content after WGD in cancer cells, resulting in 4N cells that differ in size. We find that small size is associated with enhanced cell fitness, mitotic fidelity, and tumorigenicity in 4N cancer cells and with poor patient survival in WGD-positive human cancers. Overall, these results suggest that cell and nuclear size contribute to the tumorigenic potential of 4N cancer cells and could be an important prognostic marker in human tumors that undergo WGD.

**Statement of Significance:** We report that WGD generates tetraploid cancer cells that vary in size, with larger cells displaying high chromosomal instability and smaller cells exhibiting high fitness and tumorigenicity. Furthermore, WGD status and cancer cell nuclear size in human tumors correlated with patient survival, demonstrating the clinical relevance of this association.

## Introduction

Whole genome doubling (WGD) events have played a prominent role in the evolution of species across the tree of life (1). In humans and other multicellular organisms, some cell types undergo WGD as a part of normal tissue development, but this often results in permanent exit from the cell cycle (2). When it emerges as a result of a non-programmed event, WGD can contribute to aging and disease, such as cancer (2). WGD is one of the most common genomic alterations in cancer and is associated with drug resistance, metastasis, and poor clinical outcome in several tumor types (3-5). The genetic redundancy afforded by WGD is thought to attenuate the deleterious effects of gene mutations and chromosome missegregation, thereby enabling the propagation of genomic and functional diversity that promote cancer evolution (6,7). This is supported by several lines of evidence showing that WGD leads to the accumulation of aneuploidy (8,9), or an abnormal chromosome number, and is sufficient to induce tumorigenesis in mice (10).

In addition to its impact on genome content, WGD also results in several non-genetic changes (11,12), including an increase in cell and nuclear size (13,14). Changes in cell and nuclear size can affect cell physiology by altering intracellular transport distances, cytoskeletal structure, gene expression, mitochondrial function, the surface area-to-volume ratio, and the DNA content-to-cytoplasm ratio (15-19). While normal cell types maintain their size within a specific range (20), cell and nuclear size abnormalities are common in cancer and associated with disease stage and progression (21). This indicates that the regulation of cell and nuclear size is important for cell physiology and tissue health (22).

In normal cells that undergo WGD, cell and nuclear size tend to scale proportionally with DNA content (13,23), such that tetraploid (4N) cells are roughly twice as big as their diploid (2N) progenitors. In cancer cells, however, cell and nuclear size do not necessarily correlate with, nor depend on, changes in DNA content (13,24,25). We previously found that cell and nuclear volume do not always scale with DNA content following WGD in DLD1 colorectal cancer cells, resulting in 4N cells of different sizes (13). High levels of nuclear size heterogeneity have also been observed in clinical samples from sarcomas and breast cancer metastases that underwent WGD (26,27). Although some studies have previously highlighted a link between cell size and function (17-19,22,28-30), there are no reports on how alterations in cell and nuclear size following WGD affect 4N cancer cell physiology and tumorigenic potential. We addressed this question by inducing WGD in breast and colon cancer cell lines and performing *in vitro* and *in vivo* functional assays on a panel of 4N clones characterized by small and large cell and nuclear volumes. We also explored the effects of WGD on cancer cell nuclear size, gene expression, and patient survival in human tumors using genomic, transcriptomic, and histopathology data from The Cancer Genome Atlas (TCGA). Altogether, our findings demonstrate that variations in cell and nuclear size are associated with the fitness and tumorigenic potential of cancer cells that undergo WGD.

## Results

### Cell and nuclear volume do not scale with DNA content after WGD in breast and colon cancer cells

We previously found that WGD generates 4N cells with varying cell and nuclear sizes in DLD1 colon cancer cells (13). To determine if this varying effect of WGD on cell and nuclear size is a common occurrence in cancer cells, we induced cytokinesis failure in DLD1 (colon; mutant p53), HCT116 (colon; functional p53), and CAL51 (breast; functional p53) cells and derived single cell clones using limiting dilution (Fig. 1A). These cell lines were selected because (i) they are near-diploid (2N), as opposed to being hyper-diploid or highly aneuploid like many other cancer cell lines, (ii) their p53 status varies, which is important as p53 is commonly mutated prior to WGD in tumors (3), and (iii) WGD is prevalent in these cancer types (3-5). We then screened isolated clones by determining cell ploidy and analyzing nuclear volume in cells that expressed the cell cycle marker geminin. Geminin is expressed after the G1/S transition, a major cell size checkpoint (31,32), and is degraded during mitosis (33). Using this approach, we identified 4N clones from all three cancer cell lines that differed substantially in size but not ploidy (Fig. 1 B-H and Supplementary Fig. 1A). Other studies in mammalian cells have shown that WGD often leads to a proportional (i.e., ∼2-fold) increase in cell and nuclear size (14,18,34-36). Consistent with this, we found that nuclear volume approximately doubled, or scaled with the change in DNA content, in certain 4N clones (hereby denoted as large or L clones) compared to the parental cell lines (Fig. 1B-E). In other 4N clones (hereby denoted as small or S clones), however, nuclear volume did not scale with DNA content and only increased by 40-70% compared to 2N cells (Fig. 1B-E). For a 4N clone to be defined as S, we required that nuclear volume be ≥20% smaller than the L 4N clones (note that only one 4N clone from CAL51 met this criterion despite multiple attempts). To confirm that the size differences in the S and L 4N clones are not due to defects in cell cycle progression, we measured nuclear volume in cells positive for histone H3 phosphorylation, a marker of mitotic entry (37). In line with the nuclear volume measurements in geminin-positive cells, the L 4N clones were approximately twice the size of the 2N parental cells, while the S 4N clones were 20-25% smaller than the L 4N clones (Supplementary Fig. 1B-E).

**Figure 1.**
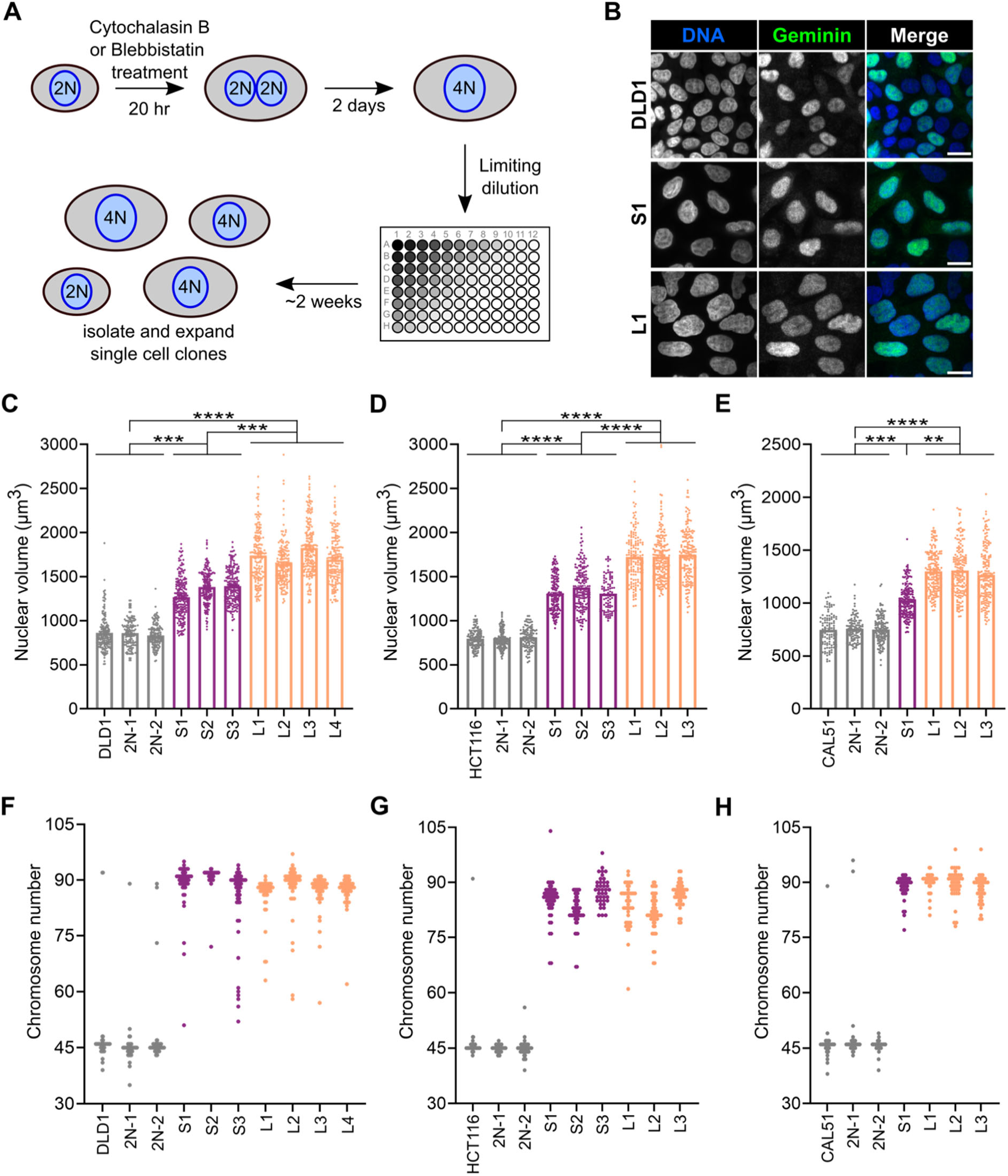
Nuclear volume does not scale with DNA content after WGD in breast and colon cancer cells. **(A)** Schematic depicting the experimental approach used to derive 4N clones. **(B)** Representative images from DLD1 2N (top), 4N S1 (middle), and 4N L1 (bottom) used for nuclear volume measurements. All scale bars = 20 μm. **(C-E)** Nuclear volume measurements in cells expressing geminin (green nuclei in panel B), a marker used to identify cells that have progressed through the G1/S transition of the cell cycle, from parental cells and 2N clones (grey), S 4N clones (purple), and L 4N clones (orange) from DLD-1, HCT-116, and CAL51 cell lines. Data are reported as mean ± SEM with individual points from three independent experiments (n ≥ 50 nuclei per experiment). ** = p<0.01, *** = p<0.001, **** = p<0.0001. **(F-H)** Quantification of chromosome numbers in metaphase spreads to determine ploidy in 2N, S 4N, and L 4N clones (n ≥ 30).

To determine if cell volume scaled with nuclear volume in the S and L 4N clones, we measured cell and nuclear volumes from individual cells in an asynchronous population. Consistent with the nuclear volume measurements (Fig. 1B-E, Supplementary Fig. 1), cell volume was significantly larger in the L 4N clones compared to the S 4N clones and 2N cells (Supplementary Fig. 2). Cell volume tended to scale with nuclear volume after WGD, and no differences in the N:C ratio were found between DLD1 and CAL51 S and L 4N clones (Supplementary Fig. 2). Despite the differences in cell and nuclear volume, the S and L 4N clones maintained near-4N genomes, with modal chromosome numbers ranging from 88-92 in DLD1 and CAL51 and 81-88 in HCT116 (Fig. 1F-H). We also determined the karyotypes of the S and L 4N clones using multicolor fluorescence *in situ* hybridization (mFISH). Consistent with our previous study (9), whole chromosome aneuploidy tended to be higher in the 4N clones compared to the 2N parental cells, and whole chromosome losses outnumbered gains in most of the 4N clones (Supplementary Figs. 3-5). While some of the 4N clones had acquired various clonal chromosome copy number alterations after WGD (e.g., DLD1-S1, S3, L4; HCT-S3, L1, L3; CAL51-S1), none of the aneuploidies were shared across all the S and/or L 4N clones (Supplementary Figs. 3-5). Therefore, the variations in cell and nuclear volume in the S and L 4N clones do not result from differences in DNA content or recurrent chromosome copy number alterations. Both chromosome numbers and nuclear volume in 2N clones (i.e., cells that did not divide during drug treatment) remained consistent with the parental population (Fig. 1B-H).

To identify transcriptional alterations associated with cell and nuclear size after WGD, we performed RNA sequencing and compared gene expression in the S and L 4N clones. Consistent with the differences in cell and nuclear size, several Gene Ontology (GO) terms associated with signaling pathways involved in cell growth and size regulation (e.g., MAPK, PI3K, ERK, IGF1R, PDGFR) were enriched among the upregulated genes in the L compared to S 4N clones across all cell lines, but most prominently in DLD1 (Supplementary Fig. 6A-C and Supplementary Table 1). There were also several cell line-specific transcriptional signatures associated with rRNA processing, mitochondrial gene expression, cell-cell adhesion, and response to chemicals in the L 4N clones (Supplementary Fig. 6A-C and Supplementary Table 1). Few pathways were found to be upregulated in the S compared to L 4N clones, except for in CAL51. GO terms associated with development and differentiation processes as well as heart contraction and cardiac conduction, which included genes encoding for plasma and mitochondrial membrane proteins involved in calcium ion transport and signaling (e.g., CACNG6, CACNG7, CACNG8, ITPR3), were enriched in the CAL51 S compared to L 4N clones (Supplementary Fig. 6D and Supplementary Table 2). In general, these findings suggest that the regulation of cell growth pathways may differ in the S and L 4N clones and contribute to the variations in cell and nuclear size.

Altogether, our results across multiple cancer cell lines show that cell and nuclear volume do not always scale with genome size upon WGD, independent of cancer type, p53 status, and chromosome copy number alterations, resulting in 4N cells with different sizes and transcriptional signatures.

### The small 4N clones display increased cell fitness compared to the large 4N clones

We next investigated whether cell and nuclear size differences after WGD affected cellular phenotypes commonly associated with transformation, such as proliferation, invasiveness, and anchorage-independent growth. Despite the oncogenic effects of WGD, 4N cells often proliferate more slowly than the 2N cells from which they are derived (8,14,38,39). To assess proliferation rate, we measured the doubling times of the S and L 4N clones and parental 2N cells. We found that the L 4N clones had longer doubling times compared to most of the S 4N clones across all cell lines (Fig. 2A-C). The L 4N clones also proliferated at a slower rate than the 2N cells in DLD1 and HCT116 (Fig. 2A-B); in CAL51, the S 4N clone proliferated at a faster rate than both the L 4N clones and the 2N parental cells (Fig. 2C). Despite the differences in size and ploidy, the CAL51 2N cells and L 4N clones as well as the DLD1 and HCT116 2N cells and S 4N clones had comparable proliferation rates (Fig. 2A-C). These findings show that WGD does not always reduce proliferation in cancer cells, but also that proliferation rate tends to decrease as cell and nuclear size increase in the 4N clones, consistent with reports in other systems (17,18,40,41).

**Figure 2.**
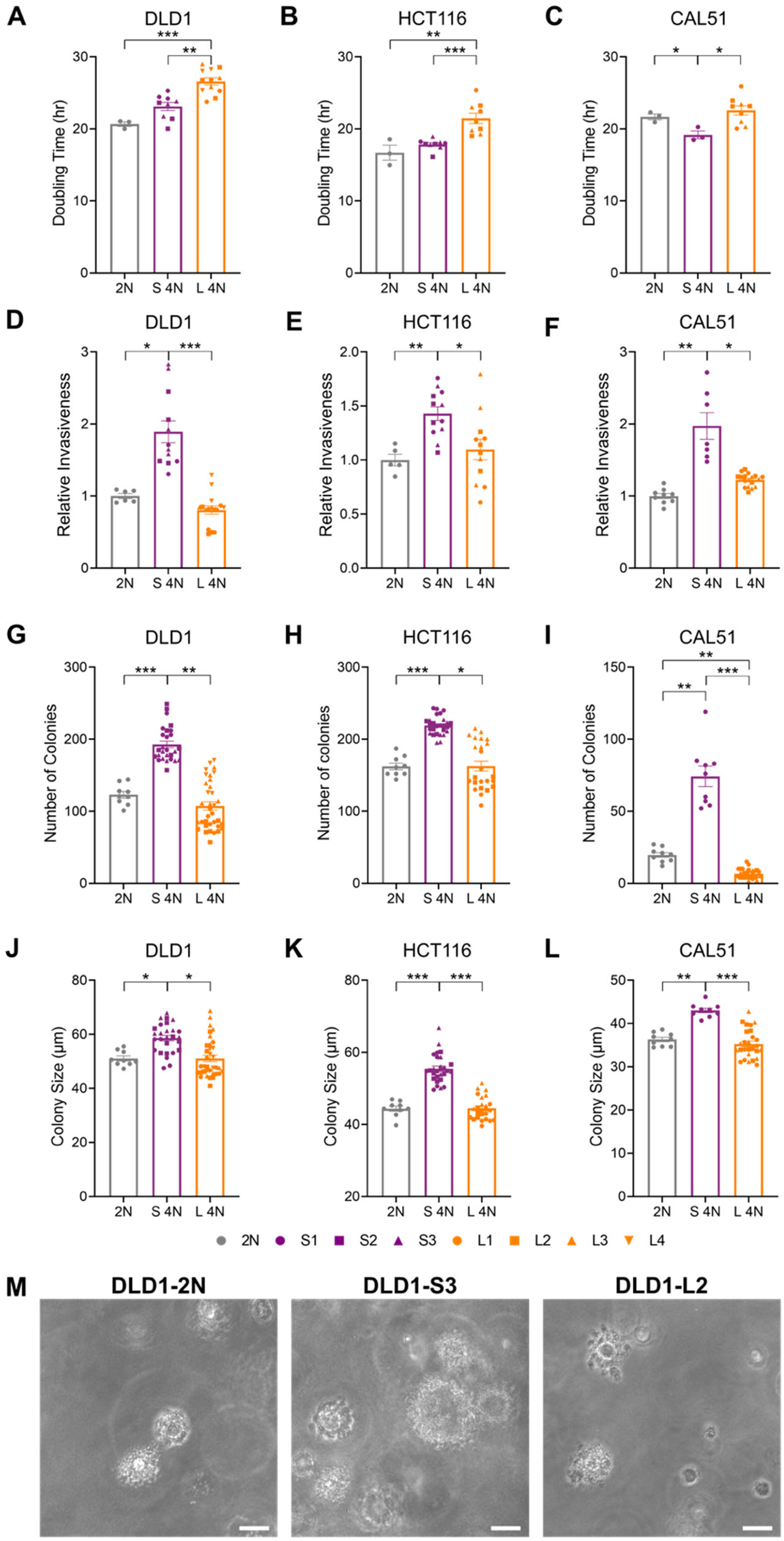
The small 4N clones display increased cell fitness compared to the large 4N clones. Quantification of **(A-C)** doubling times, **(D-F)** cellular invasion, and the **(G-I)** number and **(J-L)** size of colonies after 3 weeks in soft agar. Bar graphs report mean ± SEM for each group. Individual data points show the values of experimental replicates (n = 3) in **(A-C)** and technical replicates from **(D-F)** two and **(G-L)** three independent experiments. **(M)** Representative images of soft agar colonies from DLD1 2N (left), S3 (middle), and L2 (right). All scale bars = 30 μm. * = p<0.05, ** = p<0.01, *** = p<0.001.

To assess invasiveness, we used a basement membrane cell invasion assay and found that the S 4N clones were more invasive than both the 2N parental cells and the L 4N clones in all three cell lines (Fig. 2D-F). The L 4N clones and 2N cells displayed similar invasion efficiency, suggesting that WGD does not inherently result in increased invasiveness. Finally, we assessed anchorage-independence by growing cells in soft agar for 3 weeks. We found that the S 4N clones formed more and larger colonies than both the L 4N clones and the parental cells (Fig. 2G-M). Collectively, these *in vitro* phenotypic assays demonstrate that the level of cell fitness conferred by WGD varied, with the S 4N clones displaying higher fitness than both the L 4N clones and the parental 2N cells.

Since DLD1-S2 remained tetraploid (most cells had 92XXYY karyotypes) and had low levels of aneuploidy (Figure 1F and Supplementary Fig. 3C), we next wanted to determine if manipulating cell and nuclear size would impact the fitness of DLD1-S2 cells. To increase cell and nuclear size, we exposed DLD1-S2 cells to Palbociclib—a CDK4/6 inhibitor that arrests cells in G1 and leads to excessive cell growth (17,19,31,42-44)—for 48 hr and then plated cells at limiting dilution to isolate single cell clones that remained larger than the parental cells in the absence of the drug. We were able to isolate two clones (DLD1-S2-LV1 and LV2) characterized by a nuclear volume that was 20% larger than the parental DLD1-S2 cells (Supplementary Fig. 7A-B), while also remaining near-4N (Supplementary Fig. 7C). Consistent with our findings in the L 4N clones, we found that DLD1-S2-LV1 and LV2 were less proliferative, less invasive, and formed significantly fewer and smaller colonies in soft agar compared to the parental DLD1-S2 cells (Supplementary Fig. 7D-F). These findings suggest that cell fitness decreases as cell and nuclear size increase in 4N cancer cells.

### The large 4N clones have increased chromosomal instability compared to the small 4N clones

We next wanted to explore the possible underlying causes of the reduced fitness of the L 4N clones compared to the S 4N clones. Previous studies have shown that WGD promotes chromosomal instability (CIN) and aneuploidy (4,8,9). While CIN and aneuploidy can lead to beneficial functional adaptations, high levels of chromosome missegregation are most often detrimental to cell growth and fitness in yeast and mammalian cells (45,46). Therefore, we investigated whether the differences in cell fitness in the 4N clones could be caused by differences in the rates of chromosome segregation errors. First, we used the mFISH data to determine the levels of karyotypic variation in the S and L 4N clones by calculating the number of non-clonal (whole and structural chromosome) aneuploidies per cell. Non-clonal aneuploidy was highest in the L 4N clones (Supplementary Fig. 8A-C), suggesting that they are more prone to chromosome missegregation than the S 4N clones and 2N cells. Our prior work showed that L 4N DLD1 displayed higher rates of misaligned chromosomes compared to S 4N DLD1 clones (13), a defect that can lead to chromosome missegregation when unresolved prior to anaphase onset (47,48). Consistent with this, we found that chromosome misalignment was highest in the L 4N clones across all cell lines (Fig. 3A-D). Next, we quantified the rates of anaphase lagging chromosomes, a chromosome segregation error often observed in chromosomally unstable cancer cells and tumors (49,50), in the S and L 4N clones. The incidence of lagging chromosomes was approximately 2-fold and 4-fold higher in the L 4N clones compared to the S 4N clones and 2N cells, respectively (Fig. 3E-H). While the 2-fold increase in lagging chromosomes in the S 4N clones compared to the 2N cells could simply be due to the doubling of chromosome numbers as suggested previously (4), this cannot explain the high rates of lagging chromosomes in the L 4N clones. We also confirmed that there were no differences in the fraction of mitotic cells with extra centrosomes, which can lead to chromosome missegregation (51,52), in the S and L 4N clones and 2N parental cells (Supplementary Fig. 8D-F). Lastly, we quantified the frequency of micronuclei in interphase cells, as they can form from anaphase lagging chromosomes upon mitotic exit (53). Consistent with the observed rates of anaphase lagging chromosomes, the fraction of interphase cells with micronuclei was higher in the L 4N clones compared to the S 4N clones and the parental cells (Fig. 3I-K). Overall, these data show that the L 4N clones display higher rates of chromosome missegregation than the S 4N clones and the 2N parental cells, which may contribute to the lower fitness of the L 4N clones.

**Figure 3.**
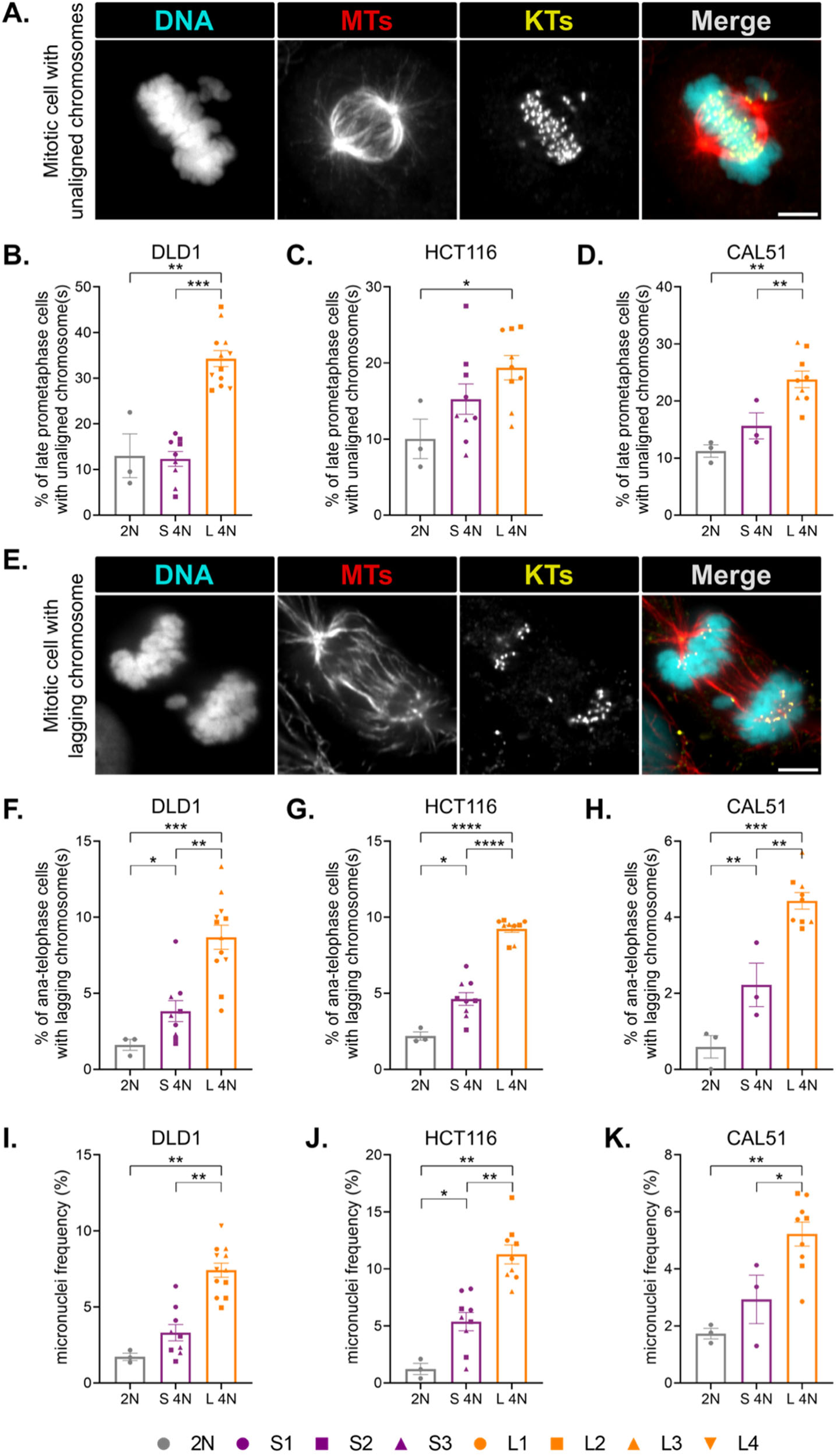
The large 4N clones are more prone to chromosome misalignment and missegregation compared to the small 4N clones. **(A)** Representative image of a mitotic cell with a distinct metaphase plate and unaligned chromosomes near one of the spindle poles. **(B-D)** Quantification of mitotic cells with unaligned chromosomes. **(E)** Representative image of a mitotic cell with a lagging chromosome. **(F-H)** Quantification of ana-telophase cells with lagging chromosome(s). **(I-K)** Quantification of the fraction of interphase cells with micronuclei. Data depicted as mean ± SEM for each group with individual points showing the means of experimental replicates (n = 3) from each 2N cell line and 4N clone. A minimum of 100 and 500 cells were analyzed per experiment in (B-D, F-H) and (I-K), respectively. All scale bars = 5 μm. * = p<0.05, ** = p<0.01, *** = p<0.001, **** = p<0.0001.

### Protein synthesis and mitochondrial content tend to scale with ploidy rather than cell and nuclear volume in the 4N clones

Previous studies found that protein synthesis rates and metabolic activity did not completely scale with size in large cells (15,17,40,41), which could further explain the reduced proliferation and fitness in our L 4N clones. Thus, we aimed to determine the relationship between biosynthesis and size in the S and L 4N clones. To do this, we first measured protein synthesis by quantifying O-propargyl-puromycin (OPP) accumulation in nascent polypeptides (Fig. 4A). Consistent with a higher demand for proteins in the 4N compared to 2N cells, nascent polypeptide abundance (or total OPP fluorescence intensity) was higher in the S and L 4N clones than in the parental 2N cells across all cell lines, although this difference did not reach statistical significance in the DLD1 L 4N clones (Fig. 4B-D). Notably, nascent protein levels did not differ between the S and L 4N clones (Fig. 4B-D), suggesting that protein synthesis may scale with cell size differently in the 4N clones. To explore this further, we examined the relationship between total OPP abundance and cell area in the 4N clones and found that OPP accumulated more rapidly in the S compared to the L 4N clones (Supplementary Fig. 9A-C). Furthermore, the concentration of nascent polypeptides (or mean OPP intensity) per cell also tended to be higher in the S compared to the L 4N clones, although the difference was only statistically significant in CAL51 (Supplementary Fig. 10A-C).

**Figure 4.**
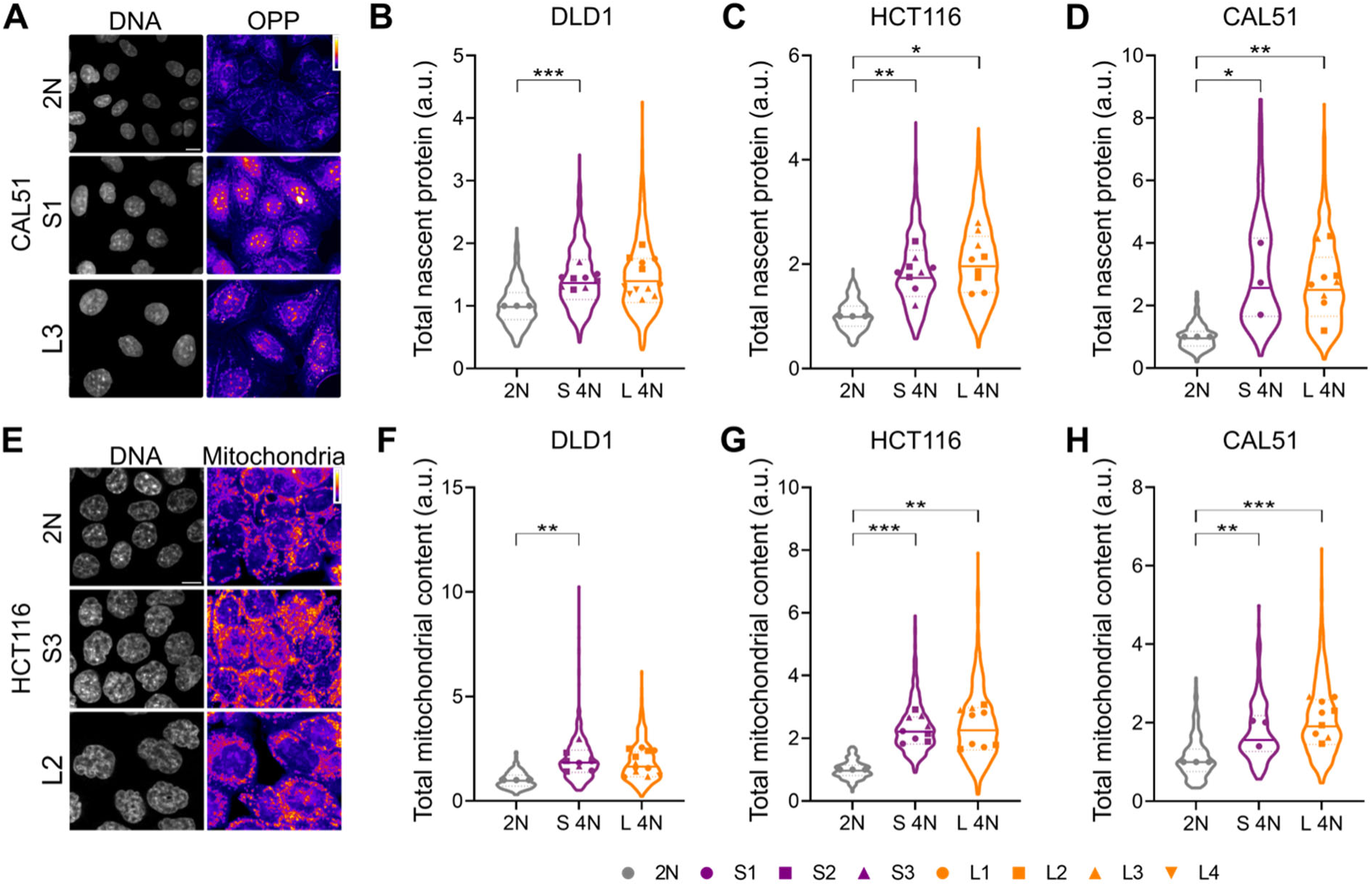
Protein synthesis and mitochondrial content do not scale with cell and nuclear size in the 4N clones. **(A)** Representative images showing OPP accumulation in nascent polypeptides in CAL51 2N (top), S1 (middle), and L2 (bottom) cells. **(B-D)** Quantification of the total OPP fluorescence intensity per cell. **(E)** Representative images of mitochondria fluorescently labeled with MitoTracker Deep Red FM in HCT116 2N (top), S3 (middle), and L2 (bottom) cells. **(F-H)** Quantification of total mitochondrial content per cell. In all graphs, individual data points depict the means of experimental replicates (n = 3) for the 2N cells and 4N clones. Violin plots display all fluorescence intensity measurements for the 2N, S 4N, and L 4N groups. Bold and dashed lines correspond to the median and quartiles, respectively. All scale bars = 10 μm. * = p<0.05, ** = p<0.01, *** = p<0.001.

Since protein synthesis rate is linked to metabolic activity (15), we next quantified the content of active mitochondria in the S and L 4N clones. Across all cell lines, total mitochondrial content per cell was higher in the 4N clones compared to the parental 2N cells but did not differ between the S and L 4N clones (Fig. 4E-H). Additionally, mitochondrial content was higher in the S 4N clones compared to the L 4N clones even when cell sizes were similar (Supplementary Fig. 9D-F), and mitochondrial content increased more rapidly as a function of size in DLD1 and CAL51 S 4N clones compared to the L 4N clones (Supplementary Fig. 9D, F). We also found that the S 4N clones had the highest mitochondrial density (Supplementary Fig. 10D-F). Therefore, consistent with protein synthesis levels, mitochondrial content increased after WGD but did not scale with cell and nuclear size in the S and L 4N clones. Taken together, these data indicate that, despite their larger size, the L 4N clones do not synthesize more proteins and have lower mitochondrial density compared to the S 4N clones. Thus, along with the high levels of CIN, the low fitness of the L 4N clones may result from biosynthetic and metabolic deficiencies associated with large cell size (15,17,40,41).

### The large 4N clones are more vulnerable to proteotoxic and oxidative stress than the small 4N clones

Due to the high rate of CIN and inability to scale protein synthesis and mitochondrial content with cell and nuclear size in the L 4N clones, we reasoned that disrupting proteostasis and metabolic function would be more costly in the L 4N clones compared to the S 4N clones. To test this, we induced proteotoxic stress using tunicamycin, a drug that induces protein misfolding, and the proteasome inhibitor bortezomib. We then measured long-term cell viability by analyzing the clonogenicity after drug exposure relative to the untreated control. The S 4N clones displayed higher tolerance to tunicamycin compared to the L 4N clones across all cell lines (Fig. 5A-D) and, in some cases, also compared to the 2N cells (Fig. 5B, D). We observed a similar trend with bortezomib (Fig. 5E-H). Since tunicamycin and bortezomib can also cause oxidative stress (54,55), we next assessed whether the L 4N clones were more sensitive to oxidative stress than the S 4N clones by performing a clonogenic assay in cells treated with tert-butyl hydroperoxide (tBHP). We found that colony formation was higher in the S compared to the L 4N clones across a range of tBHP concentrations in all three cell lines (Fig. 5I-L), suggesting that L 4N clones are more vulnerable to oxidative stress than the S 4N clones. Overall, these findings demonstrate that the L 4N clones have lower tolerance to proteotoxic and oxidative stress compared to the S 4N clones, which could be linked to the high rate of chromosome missegregation (Fig. 3) and the inability of L 4N clones to scale biosynthesis with cell and nuclear size (Fig. 4).

**Figure 5.**
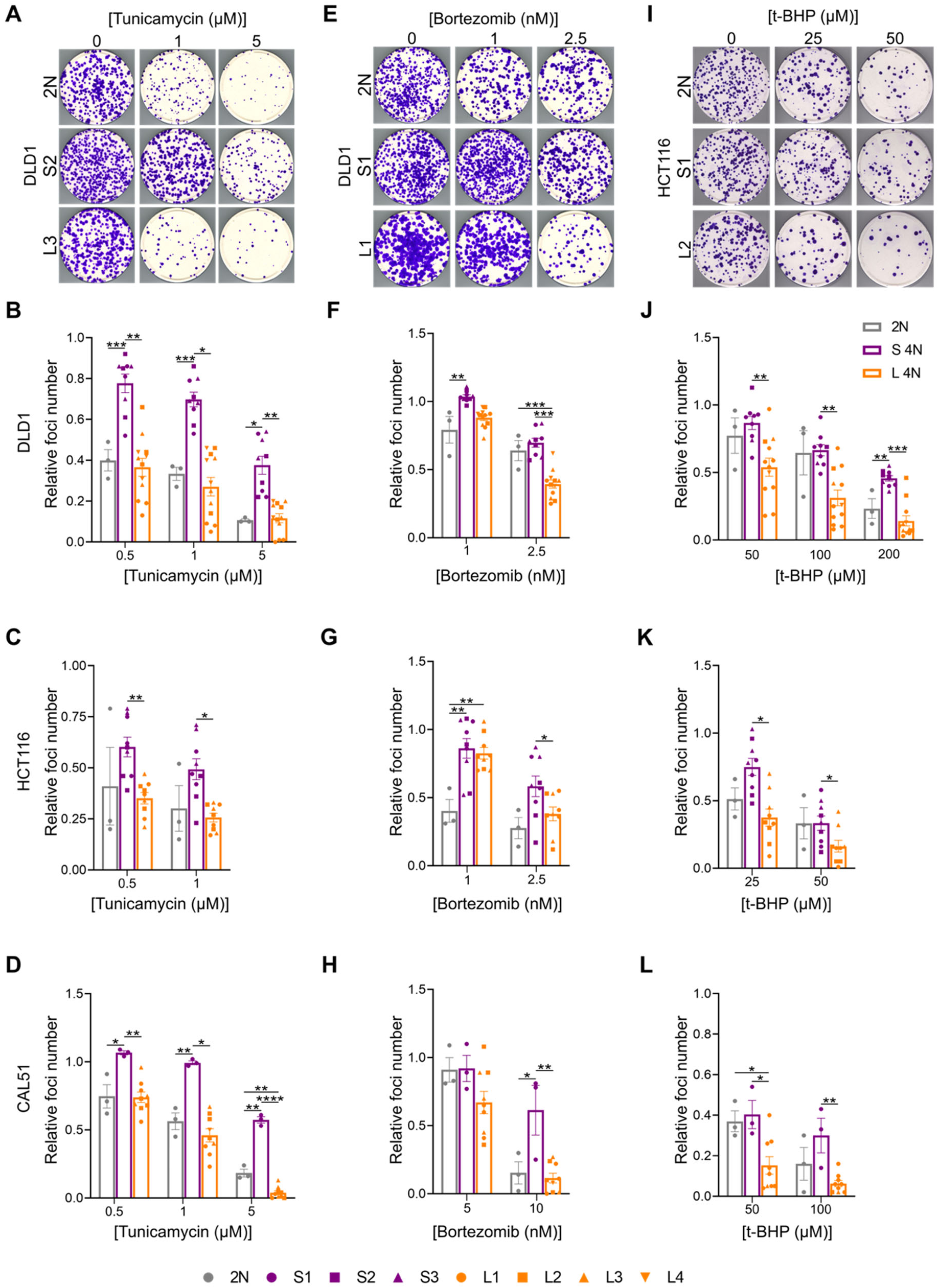
The large 4N clones are more vulnerable to proteotoxic and oxidative stress than the small 4N clones. **(A)** Representative images of foci formed by DLD1 2N (top), S2 (middle), and L3 (bottom) after exposure to DMSO or tunicamycin. **(B-D)** Quantification of relative number of foci in treatment group compared to control group for each 2N cell line and 4N clone. **(E)** Representative images of foci formed byDLD1 2N (top), S1 (middle), and L1 (bottom) after exposure to DMSO or bortezomib. **(F-H)** Quantification of relative number of foci in treatment group compared to control group for each 2N cell line and 4N clone. **(I)** Representative images of foci formed by DLD1 2N (top), S1 (middle), and L1 (bottom) after exposure to t-BHP. **(J-L)** Quantification of relative number of foci in treatment group compared to control group for each 2N cell line and 4N clone. All graphs report data as mean ± SEM for each group with individual points corresponding to the value for each independent experiment (n = 3) from each 2N cell line and 4N clone. * = p<0.05, ** = p<0.01, *** = p<0.001.

### The small 4N clones are more tumorigenic than the large 4N clones

All the functional assays performed in cell culture indicated that the L 4N clones have lower fitness than the S 4N clones (Fig. 2), suggesting that they may also be less tumorigenic. To assess this, we injected cells into the flanks of nude mice and measured tumor growth. Consistent with the fitness differences in the *in vitro* assays, the S 4N clones showed higher tumorigenicity in nude mice compared to the L 4N clones, while displaying similar tumorigenic potential as the parental 2N cells (Fig. 6A-D). To determine if nuclear size changed during *in vivo* tumor formation, we isolated cancer cells from the xenograft tumors that reached a suitable size for cell recovery and measured nuclear volume in the S and L 4N clones. Across all cell lines, nuclear volume in the post-xenograft cells did not change compared to the pre-xenograft cells in the S 4N clones and 2N cells (Fig. 6E, G, I), indicating that nuclear volume in these cells is stable after recovery from the xenograft tumors. In the DLD1 and HCT116 L 4N clones, we found that there was a modest decrease in nuclear volume in the post-xenograft cells compared to the pre-xenograft cells (Fig. 6E, G). Quantification of chromosome numbers from the post-xenograft cell populations showed that many cells remained near-4N and there were similar levels of chromosome number heterogeneity in the DLD1 and HCT116 S and L 4N clones, while chromosome numbers in the CAL51-L3 post-xenograft cells (note that the CAL51-L3 clone was the only L 4N clone that formed tumors) became highly variable and trended towards near-triploid (3N) karyotypes (Fig. 6F, H, I). This indicates that although chromosome numbers become more heterogeneous during tumor formation, chromosome loss does not necessarily result in decreased nuclear size in 4N cancer cells. Altogether, these findings demonstrate that the S 4N clones are more tumorigenic than the L 4N clones, suggesting that reducing cell and nuclear size could be favorable for tetraploidy-mediated tumor development.

**Figure 6.**
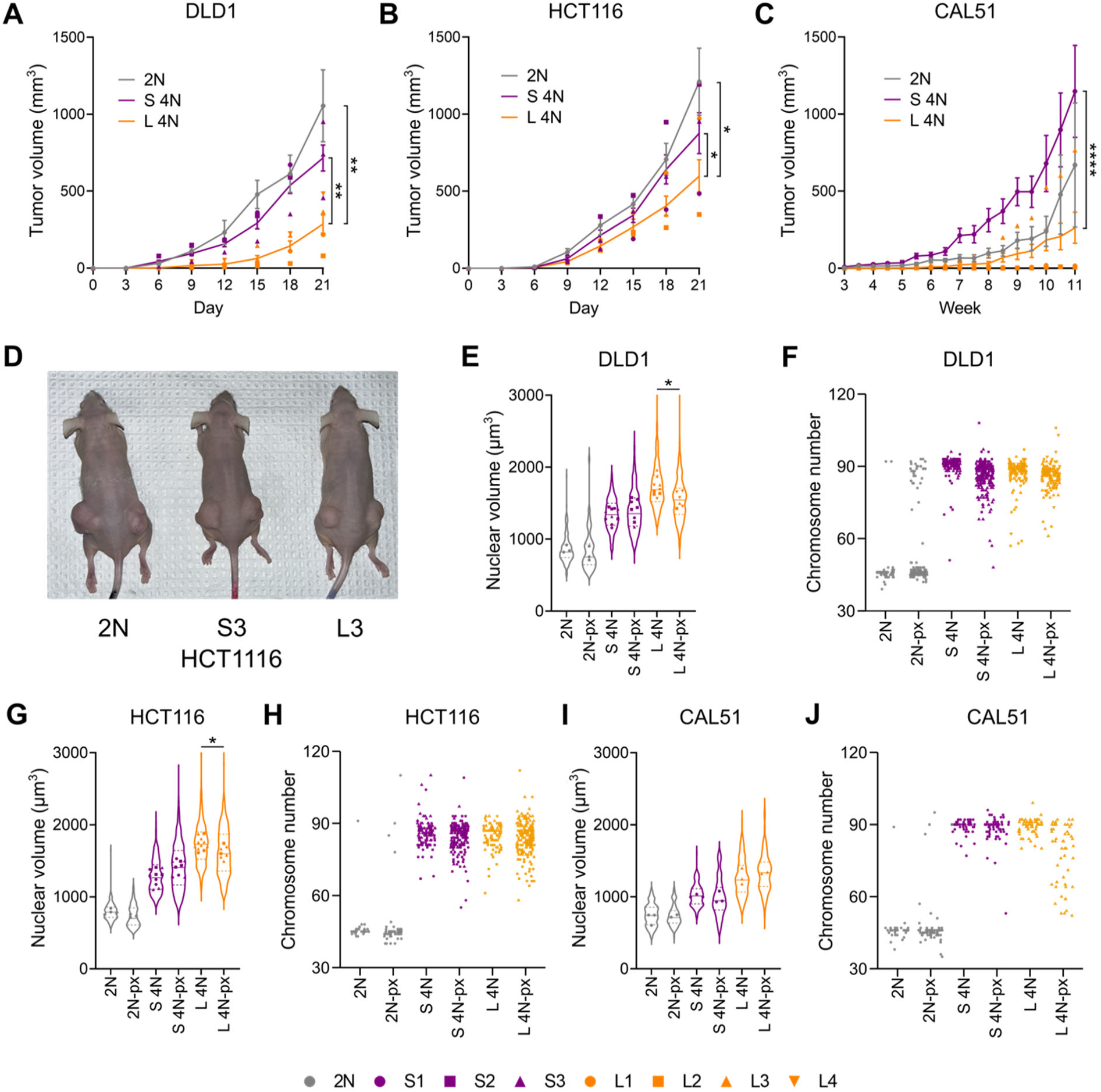
The small 4N clones are more tumorigenic than the large 4N clones. **(A-D)** Quantification and representative images of xenograft tumor growth in nude mice. 2N cells, S 4N clones, and L 4N clones from each cell line were subcutaneously injected into the flanks of nude mice (n = 8-10 xenograft tumors per clone). For the 2N group, the line and error bars show the median ± SEM at each time point. For the S and L 4N groups, individual points correspond to the median tumor volume for each 4N clone, which were used to calculate the mean ± SEM for each group and are depicted by the line and error bars. Significance was assessed by comparing the slopes of a simple linear regression. **(E, G, I)** Nuclear volume measurements in geminin-positive cells from the 2N, S 4N, and L 4N cells recovered from xenograft tumors (denoted by “px” subfix) reported alongside pre-xenograft measurements. In pre-xenograft groups, individual data points for 2N cell lines and 4N clones display the average nuclear volume from independent experimental replicates reported in Fig. 1 (n = 3-4 per 2N cell line or 4N clone). In post-xenograft (px) groups, individual data points correspond to mean nuclear volume for cells recovered from different tumors (n = 3-4). Violin plots display all nuclear volume measurements for the 2N, S 4N, and L 4N groups. Bold and dashed lines correspond to the median and quartiles, respectively. **(F, H, J)** Quantification of chromosome numbers in metaphase spreads from the 2N, S 4N, and L 4N cells before and after recovery from xenograft tumors. * = p<0.05, ** = p<0.01, **** = p<0.0001.

### Nuclear size varies among WGD-positive human tumors and is associated with patient survival

WGD, one of the most common genomic alterations in cancer, is associated with poor survival and metastasis in several cancer types (3,4), making WGD status an important prognostic factor. However, our cell line data suggest that size should also be considered, given the effect on cell fitness and tumorigenic potential. Several studies have found that genomic and molecular data from cancer cell lines can recapitulate observations in human tumors and provide important insights into cancer progression and potential therapeutic targets (9,56-59). Therefore, we next aimed to determine whether size is associated with patient survival in human WGD-positive (WGD+) cancers. To measure nuclear size in patient samples, we analyzed histopathology images from The Cancer Genome Atlas (TCGA) using the automated computational pathology toolkit TIA Toolbox (60). We used TIA Toolbox features to identify cancerous regions within tissue biopsies and then perform nuclear segmentation and classification to measure nuclear area for different cell types (e.g., neoplastic, immune, etc.) in human tumors (see Methods) (60). Previous studies have characterized the prevalence of WGD in the TCGA database (57,61,62), which allowed us to group patient samples by WGD status. Using this approach, we analyzed nuclear area in more than 17 million neoplastic cells from WGD-negative (WGD-) and WGD+ carcinomas (1095 patient samples in total) from 6 tumor types in which WGD is prevalent (∼40% or higher), including bladder urothelial carcinoma (BLCA), breast invasive carcinoma (BRCA), colon adenocarcinoma and rectum adenocarcinoma (CRAD), esophageal adenocarcinoma (EAC), lung adenocarcinoma (LUAD), and stomach adenocarcinoma (STAD). To be consistent with our *in vitro* model, we restricted the analysis of WGD-cancers to those with a low aneuploidy score (≤3 chromosomal alterations)—except in EAC as this was not possible due to the low number of samples. We found that neoplastic cell nuclear area was higher in WGD+ compared to WGD− cancers in all tumor types except EAC (Fig. 7A), while immune cell nuclear size did not differ between WGD- and WGD+ cancers, as expected (Fig. 7B). Consistent with our cell line model, there was also a high degree of variation in neoplastic cell nuclear size between the WGD+ cancers within each tumor type (Fig. 7A), enabling us to categorize the WGD+ samples into groups based on median neoplastic cell nuclear size (Fig. 7C-D). Across all tumor types analyzed, the large WGD+ group, or the top third of samples, had a median neoplastic cell nuclear size that was approximately 30% higher than the small WGD+ group, or the bottom third (Fig. 7C). In all tumor types except LUAD, aneuploidy score and neoplastic cell nuclear size were not correlated in the WGD+ cancers (Supplementary Fig. 11A-F). Similarly, there was no correlation between neoplastic cell nuclear size and tumor ploidy in any tumor type except CRAD (Supplementary Fig. 11G-L). These findings suggest that the variation in nuclear size among WGD+ cancers is not due to the accumulation of aneuploidy or loss of genomic content.

**Figure 7.**
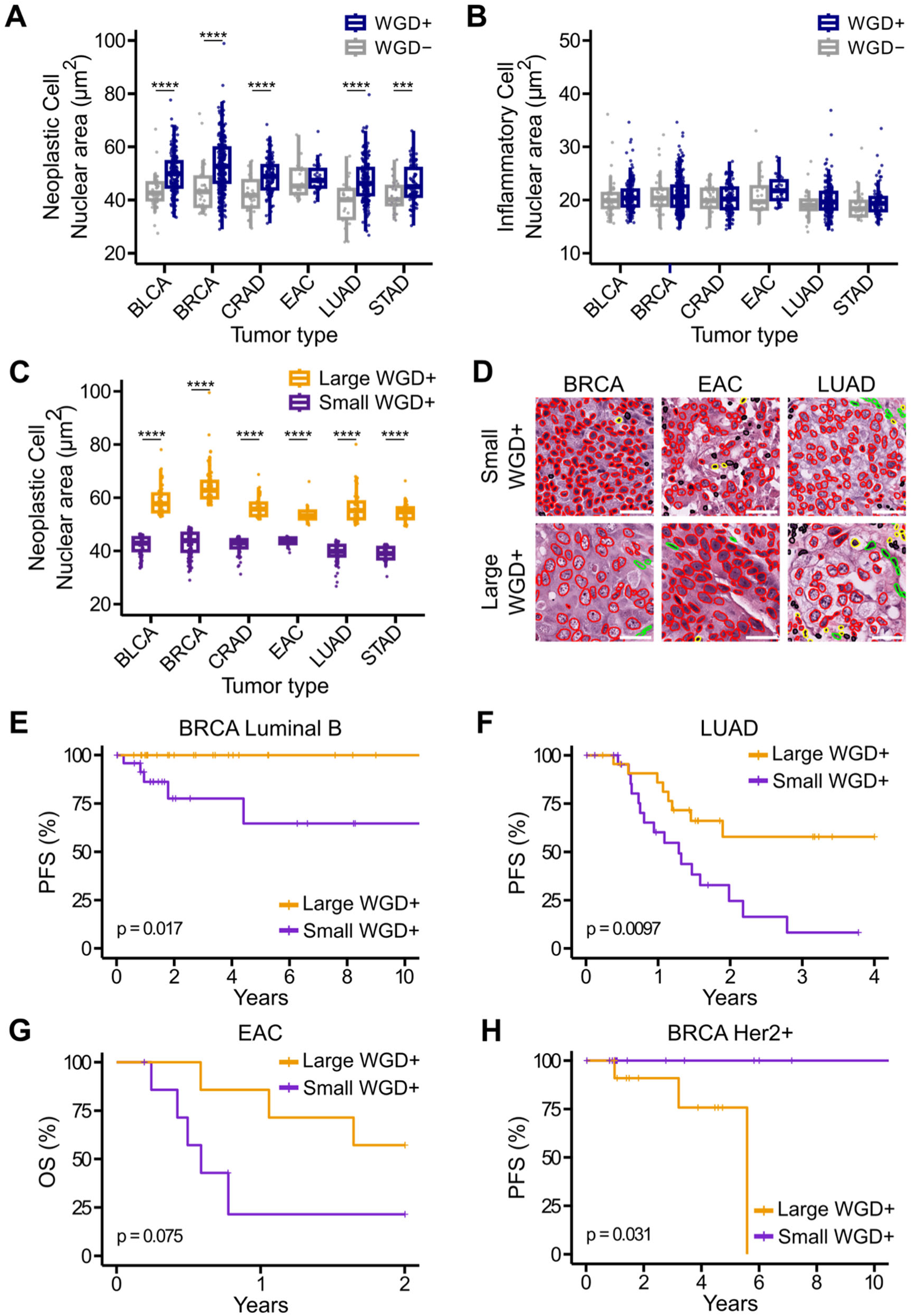
Nuclear size after WGD varies between tumor types and is associated with patient survival. Nuclear area measurements from **(A)** neoplastic and **(B)** inflammatory cells in WGD- and WGD+ cancer from six tumor types displaying high rates of WGD and for which genomics and histopathology data are available, including bladder urothelial carcinoma (BLCA), breast invasive carcinoma (BRCA), colon adenocarcinoma and rectum adenocarcinoma (CRAD), esophageal adenocarcinoma (EAC), lung adenocarcinoma (LUAD), and stomach adenocarcinoma (STAD). **(C)** Neoplastic cell nuclear area measurements in small (purple) and large (orange) WGD+ cancers from each tumor type. For all box and whisker plots, individual points depict median neoplastic cell nuclear size from each patient sample analyzed. Box plots show the interquartile range (IQR) with a line at the median and whiskers extending to smallest and largest non-outlier values (defined as those within 1.5xIQR). A two-sample t-test was used to compare WGD- and WGD+ or small and large WGD+ groups for each tumor type. **(D)** Representative images from WGD+ patient samples with small (top) and large (bottom) cancer cell nuclei in different tumor types. Nuclei from neoplastic, immune/inflammatory, connective, and dead cells are outlined in red, yellow, green, and black, respectively. All scale bars = 40 μm. **(E-H)** Kaplan-Meier survival curves in WGD+ **(E)** BRCA Luminal B, **(F)** LUAD, **(G)** EAC, and **(H)** BRCA Her2+ tumors with small (purple) and large (orange) cancer cell nuclear size. Survival curves were compared using a log-rank test. *** = p<0.001, **** = p<0.0001.

To determine if there were any transcriptional signatures associated with neoplastic cell nuclear size, we used the TCGA RNA-seq data to perform gene set enrichment analysis (GSEA) on the differentially expressed genes in the small and large WGD+ groups from each tumor type. This identified transcriptional differences in pathways associated with the cell cycle, cell-cell adhesion, development and differentiation, DNA synthesis and repair, biosynthetic and metabolic processes, ion transport, oxidative stress, and immune response between the small and large WGD+ cancers in BLCA, BRCA, LUAD, and STAD (Supplementary Fig. 12 and Supplementary Table 3). In the BLCA, BRCA, and LUAD small WGD+ cancers, most of the enriched transcriptional signatures were in cell cycle, biosynthetic, and metabolic processes, including fatty acid and RNA metabolism as well as transcription of E2F targets (Supplementary Fig. 12A, C and Supplementary Table 3). In BLCA, BRCA, and LUAD, many of the enriched gene sets in the large WGD+ cancers were in pathways associated with immune response, such as lymphocyte activation, interferon signaling, and allograft rejection (Supplementary Fig. 12A-C and Supplementary Table 3). These results demonstrate that the transcriptional regulation of immune, biosynthetic, and metabolic pathways is associated with cancer cell nuclear size in WGD+ tumors.

Finally, we asked whether there was any association between small cancer cell nuclear size and patient survival in WGD+ cancers. Since tumor histology is associated with prognosis, we first grouped patient samples by histological subtype and then, when applicable, histological grade or tumor stage (II and III were merged into a single group, while I and IV, which make up a minor fraction of cases, were not included for survival analysis), for all tumor types except BRCAs, which were grouped by molecular subtype (e.g., Her2+, Triple Negative, etc.). We next separated the samples into small and large WGD+ groups based on median neoplastic cell nuclear size and compared progression-free survival (PFS) and overall survival (OS) in these two groups. We found that the small WGD+ group had a worse prognosis than the large WGD+ group in some tumor types. In BRCA Luminal B and LUAD, PFS was lower in the small WGD+ group compared to the large WGD+ group (Fig. 7E-F). The small WGD+ group also had lower overall survival than the large WGD+ group in EAC, although this difference was not significant (Fig. 7G). There was only a single tumor subtype, BRCA Her2+, where PFS was lower in the large compared to the small WGD+ group (Fig. 7H). We also performed multivariate analyses to account for the effects of other clinical variables (e.g., patient age, gender (when applicable), and Ki67 levels). Although the hazard ratio could not be determined for some of the analyses in the BRCA tumor subtypes due to the lack of events within either the small or large WGD+ group, small neoplastic cell nuclear area was associated with poor survival in EAC and LUAD (Supplementary Table 4). Altogether, consistent with our cell line data, the patient survival data demonstrate that small neoplastic cell nuclear size in WGD+ human cancers is associated with a poor prognosis across multiple cancer types.

## Discussion

WGD is recognized as an important event in cancer progression for its effects on genome stability (3,4). There is growing evidence, however, that non-genetic alterations associated with WGD can also alter cell physiology and contribute to cancer evolution (12). Our work sheds light on a link between variations in cell and nuclear size and the fitness of 4N cancer cells. Consistent with our earlier study (13), we found that cell and nuclear size do not always scale with genome size after WGD in colorectal and breast cancer cell lines, independent of p53 status. Functional assays revealed that the S 4N clones are more fit and tumorigenic than the L 4N clones, which can be attributed to the high rate of chromosome missegregation and inability to scale biosynthesis with size in the L 4N clones. Collectively, our findings demonstrate that cell and nuclear size are associated with fitness, CIN, and tumorigenic potential in 4N cancer cells.

Recent findings indicate that the inability of large cells to scale biosynthesis and mitochondrial activity with size can impair cell proliferation and fitness (17-19), which could explain why other studies have found that stemness and tumorigenicity increase as size decreases in cancer cells (17-19,22,28,29). There is also evidence that the proteome may not scale linearly with DNA content after WGD in yeast and cancer cells (35). Consistent with these observations, our data suggest that the low fitness and tumorigenicity of the L 4N clones stems, at least in part, from a biosynthetic deficiency. This could be especially detrimental for the fitness of large 4N cells, since the functional demand for RNAs, proteins, lipids, and other macromolecules to sustain cell growth and other basic housekeeping processes increases with cell size. Additionally, the energetic costs associated with the biosynthetic requirements in large cells could be disadvantageous in stressful or nutrient poor conditions. Nutrient availability is a major selective force during cancer evolution and may also limit the incidence of WGD in some tumor types, such as glioblastoma (63). Lower nutrient availability (compared to *in vitro* conditions) could also explain why the S 4N clones, despite their enhanced capacity for anchorage-independent growth in soft agar, were not more tumorigenic in nude mice than the parental 2N cancer cells. Finally, compared to large cells, the more compact nature of small cells could make executing cellular processes, such as nutrient uptake, molecular transport, gene expression, and protein signaling, more efficient by increasing the surface area-to-volume ratio and the DNA content-to-cytoplasm ratio.

Although several 4N clones had acquired clonal aneuploidies after WGD, we did not find any recurrent chromosome copy number alterations shared by all the S or L 4N clones that could explain the differences in size or fitness. In line with other studies (8,9), we found that WGD results in increased chromosome missegregation (e.g., in the form of anaphase lagging chromosomes). However, we also showed that the effects of WGD on mitotic fidelity differ depending on cell and nuclear size. Specifically, the L 4N cells displayed the highest rates of both misaligned chromosomes and anaphase lagging chromosomes. Our previous study demonstrated that smaller 4N cells become rounder during mitosis than larger 4N cells (13). Since mitotic rounding is biophysically favorable for spindle assembly and chromosome capture (64), inefficient cell rounding along with a larger cytoplasmic volume may hinder chromosome alignment and, in turn, promote chromosome missegregation in the L 4N clones. Similarly, large nuclear size could disrupt chromosome capture and segregation in the L 4N clones by increasing the distance required for centrosome separation during prophase or between peripheral chromosomes and the spindle poles in early prometaphase. Recent findings indicate that WGD can alter microtubule dynamics (56), and WGD+ cells depend on KIF18A to dampen chromosome movements and maintain alignment at the metaphase plate (56,57), both important to prevent chromosome missegregation. There is also evidence that cell size can affect spindle assembly checkpoint (SAC) activity and mitotic progression (13,65). Therefore, the inability to align and segregate chromosomes effectively in the L 4N clones may depend on size-specific differences in KIF18A activity, microtubule dynamics, and/or SAC function.

Despite ongoing CIN, many of the L 4N clones were unable to establish clonal aneuploidies and maintained near-4N karyotypes, suggesting that cells likely experience a fitness penalty after missegregating chromosomes and are lost from the population over time. Previous studies have shown that chromosome missegregation can promote proteotoxic and metabolic stress (45,66,67), which may negatively impact the fitness of L 4N cells. In support of this, the L 4N clones were more vulnerable than the S 4N clones to proteotoxic and oxidative insults. Furthermore, the S, but not the L, 4N clones displayed increased resistance to tunicamycin, bortezomib, and tBHP compared to the parental 2N cells, indicating that WGD does not inherently promote tolerance to proteotoxic and oxidative stress in cancer cells. These findings also suggest that WGD status and cell/nuclear size could be indicative of potential therapeutic vulnerabilities. For instance, WGD+ cancers with large nuclear size could be susceptible to drugs that disrupt proteostasis, similar to the L 4N clones and recent observations in highly aneuploid cells (67). CIN and KIF18A could be other promising targets in WGD+ tumors with large nuclear size. Consistent with our results, large cancer cell nuclear size was associated with high levels of CIN in breast cancer histopathology images (68). Because disruption of KIF18A function exacerbates CIN specifically in WGD+ and highly aneuploid cells (56,57), large 4N cells may be particularly susceptible to KIF18A inhibition. Furthermore, a recent study found that there are a number of FDA-approved compounds capable of altering cancer cell size in different directions (30), raising the possibility of combining cell size-modifying drugs with other treatments that target size-specific vulnerabilities.

Using TCGA data, we were able to separate WGD+ cancers into different groups based on neoplastic cell nuclear area and identify size-specific transcriptional differences involving biosynthetic and metabolic processes, cell cycle, DNA repair and synthesis, and ion transport among others. Additionally, there was a striking enrichment of transcriptional signatures associated with immune response pathways in the large WGD+ cancer across multiple tumor types, which could be indicative of inflammation—a condition associated with cell enlargement (22,69)—and ongoing cancer immunoediting. It will be important to determine which, if any, genes in these pathways play an active role in size regulation, their contribution to cancer evolution, and if they represent therapeutic targets in specific tumor types. We also identified several tumor types, including Luminal B BRCA, EAC, and LUAD, where the small WGD+ group had poor patient survival. In line with the fitness differences between the S and L 4N clones, this suggests that small cancer cell size after WGD may lead to enhanced fitness and tumorigenic potential in certain cancers. Therefore, stratifying tumors by WGD status and cancer cell nuclear size could be a promising prognostic strategy.

In sum, using data from cancer cell lines and human cancers, we demonstrate that variations in cell and nuclear size following WGD contribute to the fitness and tumorigenic potential of 4N cells. By simultaneously changing genome content, cell and nuclear size, and several other biophysical properties, WGD is a powerful mechanism for altering cell physiology (11,12). Nevertheless, our understanding of how the genetic and non-genetic effects of WGD are integrated in 4N cells to promote functional variation and drive cancer evolution is still incomplete. Identifying the mechanism(s) regulating cell and nuclear size following WGD in various cell types may reveal new genetic targets and size-specific dependencies that can have important prognostic and/or therapeutic value. Our study provides an important step in this direction by showing that WGD status and cancer cell nuclear size can be predictive of prognosis in certain cancers.

## Material and methods

### Cell culture and chemicals

DLD1 (ATCC CCL-221) and HCT116 (ATCC CCL-247) cells were purchased from American Type Culture Collection (ATCC; Manassas, VA) and cultured in RPMI media with ATCC modification (Thermo Fisher Scientific – Gibco, CA, USA) and McCoy’s 5A media (ATCC), respectively. CAL51 cells were generously provided by Dr. Christian Zeirhut (UCL) and were cultured in high glucose DMEM (Thermo Fisher Scientific). All cells were maintained at 37°C and 5% CO_2_ in media supplemented with 10% fetal bovine serum (FBS; Thermo Fisher Scientific) and 1% anti-biotic and anti-mycotic (Thermo Fisher Scientific). Cells were checked for mycoplasma contamination every 3 weeks by DNA staining. Stock solutions for dihydrocytochalsin B (DCB; Sigma-Aldrich, St. Louis, MO), blebbistatin (Sigma-Aldrich), SYTO RNASelect green fluorescent cell stain (Thermo Fisher Scientific), Palbociclib (Sigma), MitoTracker Deep Red FM (Thermo Fisher Scientific), O-propargyl-puromycine (OPP; Vector Laboratories Inc., Newark, CA), tunicamycin (Thermo Fisher Scientific), and bortezomib (Thermo Fisher Scientific) were prepared in DMSO and diluted to final concentrations in cell culture media.

### Generation of single cell tetraploid clones

Cytokinesis failure was induced in DLD1 cells by adding 1.5 µg/mL DCB to the culture medium for 20 hr. In HCT116 and CAL51, cytokinesis failure was induced by adding 75 µM blebbistatin to the culture medium for 20 hr. Drugs were washed out four times in media. The cells were allowed to recover for 48 hr before performing single cell cloning via limiting dilution in 96-well plates. Wells containing single cells were identified and expanded into clonal cell lines for further screening. To derive large DLD1-S2 clones, cells were exposed to 500 nM Palbociclib for 48hr and allowed to recover 24 hr before plating at limiting dilutions.

### Preparation of metaphase spreads, chromosome counting, and multicolor fluorescence in situ hybridization (mFISH)

Metaphase spreads were prepared using a standard protocol as described previously (13). Metaphase spreads were dripped onto microscope slides and stained with 300 nM DAPI (Thermo Fisher Scientific) for 5 min in PBS. Images were acquired on a Nikon Eclipse Ti inverted microscope (Nikon Instruments Inc., NY, USA) equipped with ProScan automated stage (Prior Scientific, Cambridge, UK), CoolSNAP HQ2 CCD camera (Photometrics, AZ, USA), Lumen200Pro light source (Prior Scientific), and a 60X/1.4 NA Plan-Apochromatic objective (Nikon Instruments Inc.). Chromosome counting to determine cell ploidy was performed in FIJI. mFISH was performed with 24XCyte human probe cocktail (MetaSystems Group Inc., Medford, MA) according to manufacturer’s protocol as described in (9). Images were acquired with a Zeiss Axioplan 2 motorized microscope (Carl Zeiss, Inc., Oberkochen, Germany) equipped with an Ikaros4/ Isis4 system (MetaSystems) and analyzed using Isis 4 software (Metasystems).

### Immunofluorescence staining, confocal microscopy, and image analysis

For immunostaining, cells were grown on sterile glass coverslips and fixed with either 4% paraformaldehyde (PFA) for 18 min at room temperature or 100% ice-cold methanol (MeOH) for 10 min at -20°C. After PFA fixation, coverslips were washed 3 times for 5 min with phosphate buffered saline (PBS), and permeabilized with 0.2-0.5% Triton-X for 10 min before blocking with 10% boiled goat serum (Jackson ImmunoResearch Labs, West Grove, PA) in PHEM buffer (60 mM Pipes, 25 mM HEPES, 10 mM EGTA, 2 mM MgSO_4_, pH 7.0) for 1 hr. MeOH-fixed samples were washed 3 times for 5 min with PBS prior to the blocking step. The primary antibodies were used with the following dilutions and fixation procedure: rabbit anti-geminin [EPR14637] (Abcam, Waltham, MA), 1:400, PFA; mouse anti-phospho-histone H3 (Ser10) clone 3H10 (Sigma-Aldrich), 1:200, PFA; mouse anti-alpha-tubulin [DM1A] (Sigma-Aldrich), 1:500, MeOH; anti-centromere antigen (Antibodies Incorporated, Davis. CA), 1:100, MeOH; mouse anti-centrin 3 clone 3E6 (Abnova inc., Taiwan), 1:200, MeOH. The secondary antibodies were all purchased from Jackson ImmunoResearch Labs and used at the following dilutions: 1:200 AlexaFluor 488 anti-rabbit, 1:400 AlexaFluor 488 anti-mouse, 1:100 Red-X anti-human. Coverslips were counterstained with 300 nM DAPI, mounted on microscope slides in an antifade solution, and sealed with nail polish.

All images for cell and nuclear volume measurements as well as protein synthesis (OPP) and active mitochondrial content (MitoTracker Deep Red FM) quantification were acquired with a swept field confocal system (Prairie Technologies, WI, USA) on a Nikon Eclipse TE2000 inverted microscope equipeed with a 60X/1.4 NA Plan-Apochromatic lens, motorized ProScan stage (Prior Scientific), an XCITE 120Q light source (Excelitas Technologies, Waltham, MA; used to view cells prior to imaging), a CoolSNAP HQ2 CCD camera (Photometrics), an Agilent monolithic laser combiner (MLC400) controlled by a four channel acousto-optic tunable filter, and a multiband pass filter set.

All volume measurements were performed in cells fixed with PFA using FIJI as described previously (13). For nuclear volume measurements, only cells expressing geminin (Fig. 1), a marker to identify cells that have exited G1 phase of the cell cycle, and phosphorylated histone H3 (Supplementary Fig. 1) were analyzed. Paired cell and nuclear volume measurements (Supplementary Fig. 2) were collected from single cells seeded at a low density. DAPI was used to label and segment nuclei, and SYTO RNASelect was used at a concentration of 500 nM to label and segment the cell cytoplasm.

To quantify the fluorescence intensity of nascent protein abundance per cell, 20 µM OPP was added to the cell culture media for 1 hr before fixing with MeOH for 5 min at -20°C and performing click chemistry according to the manufacturer’s instructions (Vector Laboratories Inc.). To quantify mitochondrial content per cell, 200 nM MitoTracker Deep Red FM was added to cells in serum-free media for 30 min at 37°C before PFA fixation. For all OPP and MitoTracker experiments, the samples for all the 4N clones and 2N cells for each respective cell line were prepared together using identical conditions, and images were captured on the same day using identical microscope and imaging settings (e.g., exposure times, number of Z-steps, Z-step size, etc.). The images were analyzed by manually segmenting cells in FIJI to create ROIs for individual cells, which were overlaid onto a sum intensity projection of all Z stacks spanning the entire cell height and used to measure the total and mean fluorescence intensities for each cell. Background correction was performed using two identical ROIs drawn outside the cells to measure background fluorescence.

### Transcriptional profiling and pathway enrichment analysis

Total RNA was extracted using the RNeasy extraction kit (Thermo Fisher Scientific). RNA purity and integrity were assessed using gel electrophoresis and UV spectrophotometry. RNA-sequencing and pathway enrichment analysis were performed by Novogene (Sacramento, CA) using standard protocols. In brief, mRNA was purified from total RNA using poly-T oligo-attached magnetic beads and sequenced using an Illumina NovaSeq platform. Raw data were first processed using fastp software to clean the data, which were then mapped to the reference genome using Hisat2 v2.0.5. Gene expression was quantified using featureCounts v1.5.0-p3, and differential expression analysis was performed using the DESeq2 R package (1.20.0). Corrected P-value of 0.05 was set as the threshold for significant differential expression. Pathway enrichment analysis of differentially expressed genes was implemented by the clusterProfiler R package correcting for gene length bias. Terms with corrected P-value less than 0.05 were considered significant.

### Cell proliferation assay

50,000 cells were seeded per well in 6-well plates. The following day cells from triplicate wells were counted twice using a hemocytometer to determine cell number. This was repeated after 96 hr, and doubling times were determined using the following formula:

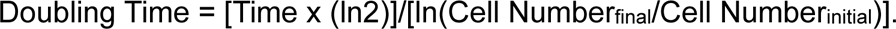

### Cell invasion assay

Invasiveness was assessed using a 96-well plate basement membrane cell invasion assay per the manufacturer’s instructions (Abcam). In brief, 40 µL of basement membrane solution were added to each well and allowed to solidify at room temperature. 50,000 cells in serum-free media were added to each well in duplicate per clone and incubated for 24 hr. Invasive cells were labeled with a fluorescent cell dye, and total fluorescence was measured using a plate reader.

### Soft agar colony formation assay

250,000 cells were seeded in 0.7% agar on top of a bottom layer with 1.0% agar in 35 mm cell culture dishes and allowed to solidify at room temperature. Cells were kept in soft agar for 3 weeks with media changes twice per week. Images spanning the entire height of the agar layer containing cells were acquired from 10 randomly selected fields of view per dish (3 dishes per experiment) using the Nikon widefield microscope described above. The major axis of each colony was measured in FIJI, and only colonies with a major axis above 30 µm were used to determine colony size and number.

### Mitotic error, micronucleus, and centriole number quantification

Cells were fixed with MeOH and immunostained, as described in an earlier section, for kinetochores and microtubules for mitotic error and micronucleus analyses, and for centrin (centriole marker) for centrosome number analysis. To quantify the frequency of chromosome misalignment, the analysis was limited to metaphase cells (i.e., cells with all chromosomes aligned at the metaphase plate) and late prometaphase cells (i.e., cells with a well-defined metaphase plate and only one or few chromosomes at or near a spindle pole). Cells in early prometaphase were excluded. Quantification of lagging chromosomes was performed in anaphase and telophase cells, while cells that had exited mitosis were excluded. Chromosomal structures in the spindle midzone without kinetochores were not scored as lagging chromosomes. All mitotic error analyses were performed blind and in triplicate (scoring at least 100 cells per experiment). To quantify the fraction of interphase cells with micronuclei, Z-stack images spanning the entire height of nuclei were acquired from randomly selected fields of view on the Nikon Eclipse Ti inverted microscope described above. Cells with and without micronuclei were counted in FIJI. Three independent experiments were performed and a minimum of 500 cells were analyzed per experiment. For centrosome analysis, the number of centrioles was determined visually in mitotic cells. Any cells with >4 centrioles were scored as having extra centrioles. Three independent experiments were performed, and 50 cells were analyzed per experiment.

### Colony formation assay

Drugs were added to the media the day after seeding 200,000 cells per well in 6-well plates. All cell lines were exposed to tunicamycin or DMSO (control) for 48 hr. For bortezomib, DLD1 and HCT116 cells were exposed to the drug or DMSO for 24 hr, and CAL51 cells were exposed to the drug or DMSO for 72 hr. For all cell lines, tert-butyl hydroperoxide (tBHP; Sigma-Aldrich) or nothing (non-treated control) was added to the media for 1 hr, then washed out three times with media, and drug-free media was added back to the cells. This procedure was repeated once per day for three days. After drug treatment, 1,000 cells were seeded into 10 cm dishes and allowed to grow for 14 days before MeOH fixation and crystal violet staining (0.5% crystal violet in 20% MeOH). Images of each dish were acquired using a flatbed scanner and colonies were counted in FIJI.

### Mouse xenograft tumor formation assay

Four-week-old female athymic Nu/J mice were purchased from The Jackson Laboratory (Bar Harbor, ME). Mice were allowed to acclimate for 2 weeks before experiments. Each flank of the mice was injected subcutaneously with 2×10^6^ cells in serum-free media diluted in Matrigel (Corning). 5 mice with injections on both flanks (n=10) were used per cell line. After the xenograft was palpable, the length and width of the tumors were measured via caliper every 3 days for DLD1 and HCT116 and twice per week for CAL51. The study was terminated when tumor size reached >15 mm in any group. To prepare cell suspensions for nuclear volume analysis, xenograft tumors were excised, minced, and digested with accutase (Sigma-Aldrich) for 30 minutes at room temperature. The cell suspension was then passed through a 70 µM pore cell filter and resuspended in normal cell culture media supplemented with 20 ng/mL human epidermal growth factor (Sigma) onto sterile glass coverslips in 35 mm cell culture dishes. To measure nuclear volume, cells were fixed with PFA 2-4 days after tumor excision and stained for geminin. Imaging and nuclear volume measurements were performed as described in the “Immunofluorescence staining, confocal microscopy, and image analysis” section.

### TCGA data analysis

TCGA data were obtained using TCGAbiolinks R package. WGD status and aneuploidy scores for TCGA samples were assigned as reported in Taylor *et al.*, Cohen-Sharir *et al*., and Quinton *et al.* (56,57,61). To measure nuclear size in TCGA diagnostic slides, the H&E whole slide images (WSIs) were processed and analyzed using TIA Toolbox, an open-source tool for tissue image analytics (60). The metadata for WSIs was used to determine the objective power and exclude WSIs that were not acquired with a 40X objective. A binary of the tissue region mask was then created, and patch extraction was performed on the masked region with a patch size of 1000×1000 pixels and a pixel size of 0.5 microns per pixel. Next, a maximum of 2,000 patches were randomly selected, normalized using the Macenko method, and classified using a pre-trained model (“resnet18-kather100k”) to identify patches extracted from tumor-rich regions of WSIs. 50 patches with tumor content were then selected at random and manually checked for quality. Patches with excessive tissue folding or pen marks were excluded from further analysis. Nuclear segmentation and classification were performed on the remaining patches using a pre-trained model (“hovernet_fast-pannuke”). Objects were grouped by classification type (e.g., neoplastic, immune, etc.), and objects with a probability score <0.9 (a measure of the likelihood a given object belongs to a certain class) or a size <10 and >250 µm^2^ were excluded. Nuclear area was calculated from the segmented object coordinates using the cv2 R package.

To compare gene expression in the small and large WGD+ groups, the DESeq function from the DESEq2 package was used with default parameters. Genes with adjusted p<0.05 were considered differentially expressed. The differentially expressed genes were ranked by log2FoldChange, and Gene set enrichment analysis (GSEA) was performed using the GSEA function from the clusterProfiler package with pvalueCutoff = 1 on HALLMARK, REACTOME, KEGG, and GO:BP gene sets (each gene set category was analyzed separately). Results with q<0.05 were considered significant.

For survival analysis, TCGA clinical data were obtained from Liu *et al*. (70). Patient samples were grouped by histological subtype and histological grade or tumor stage, excluding samples with a grade or stage of I and IV. For BRCA, patient samples were grouped by molecular subtype (e.g., Her2+, Triple Negative, etc.). The samples were then separated into small and large WGD+ groups based on median neoplastic cell nuclear area and used to compare overall survival and progression free survival. Patient survival data were censored at 2 years for ESCA as most survival events occurred within this period.

### Statistical analysis

All statistical analyses were performed in Rstudio [version 2023.03.0+386] using the following packages: tidyverse, stats, lme4, sjPlot, and survminer. General linear mixed models (GLMMs) were used to test for differences between pairs of 4N size groups and the 2N parental cells (e.g., 2N vs S 4N, 2N vs L 4N, and S vs L 4N). For statistical comparisons using GLMMs, individual clone IDs (e.g., S1, L1, S2, etc.) were categorized into two groups (small and large) based on size within each cell line. Experimental data were modeled as a function of the fixed effects variable ‘size group’ and the random effects variables ‘clone ID’ and ‘experiment ID’. A likelihood ratio test comparing models with and without the variable ‘size group’ was used to test for significant differences between pairs of the small and large 4N clone groups and the 2N group. GLMMs were used to test for significance unless specified otherwise in the figure legends. The number of replicates for each experiment is also reported in the figure legends. Significance between all group pairs (e.g., 2N vs S 4N, 2N vs L 4N, and S vs L 4N) was assessed, but only comparisons that reached statistical significance are denoted in the figures.

## Acknowledgements

We would like to thank all members of the Cimini lab and our colleagues Jing Chen, Siobhan Craige, and Silke Hauf for helpful discussions and advice. We are especially grateful to Dr. Christian Zeirhut for providing the CAL51 cells and Douglas Weidemann for help with statistical analysis. Work in the Cimini lab is supported by NIH grants R01GM140042 to D.C. and E.M.S. and 1F31CA271763-01A1 to M.B. Work in the Ben-David lab is supported by an ERC Starting Grant 945674 to U.B.-D. and a Safra Fellowship from the Edmond J. Safra Center for Bioinformatics at Tel Aviv University to R.S. We apologize to all our colleagues whose work we could not cite due to space constraints.

## Authors’ Contributions

**M. Bloomfield:** Conceptualization, data curation, formal analysis, supervision, validation, investigation, methodology, visualization, writing—original draft. **S. Huth:** Formal analysis, investigation, writing—reviewing and editing. **D. McCausland:** Formal analysis, investigation, writing—reviewing and editing. **R. Saad:** Formal analysis, investigation, methodology, visualization, writing—reviewing and editing. **N. Bano:** Methodology, writing—reviewing and editing. **T. Chau:** Methodology, writing—reviewing and editing. **M. Sweet:** Methodology, writing—reviewing and editing. **N. Baudoin:** Methodology, writing—reviewing and editing. **A. McCaffrey:** Formal analysis, investigation, writing—reviewing and editing. **K. Fluet:** Formal analysis, investigation, writing—reviewing and editing. **E.M. Schmelz:** Supervision, funding acquisition, methodology, writing—reviewing and editing. **U. Ben-David:** Supervision, funding acquisition, investigation, methodology, writing—reviewing and editing. **D. Cimini:** Conceptualization, supervision, funding acquisition, investigation, validation, methodology, writing—reviewing and editing.

---

**Supplementary Figure 1.**
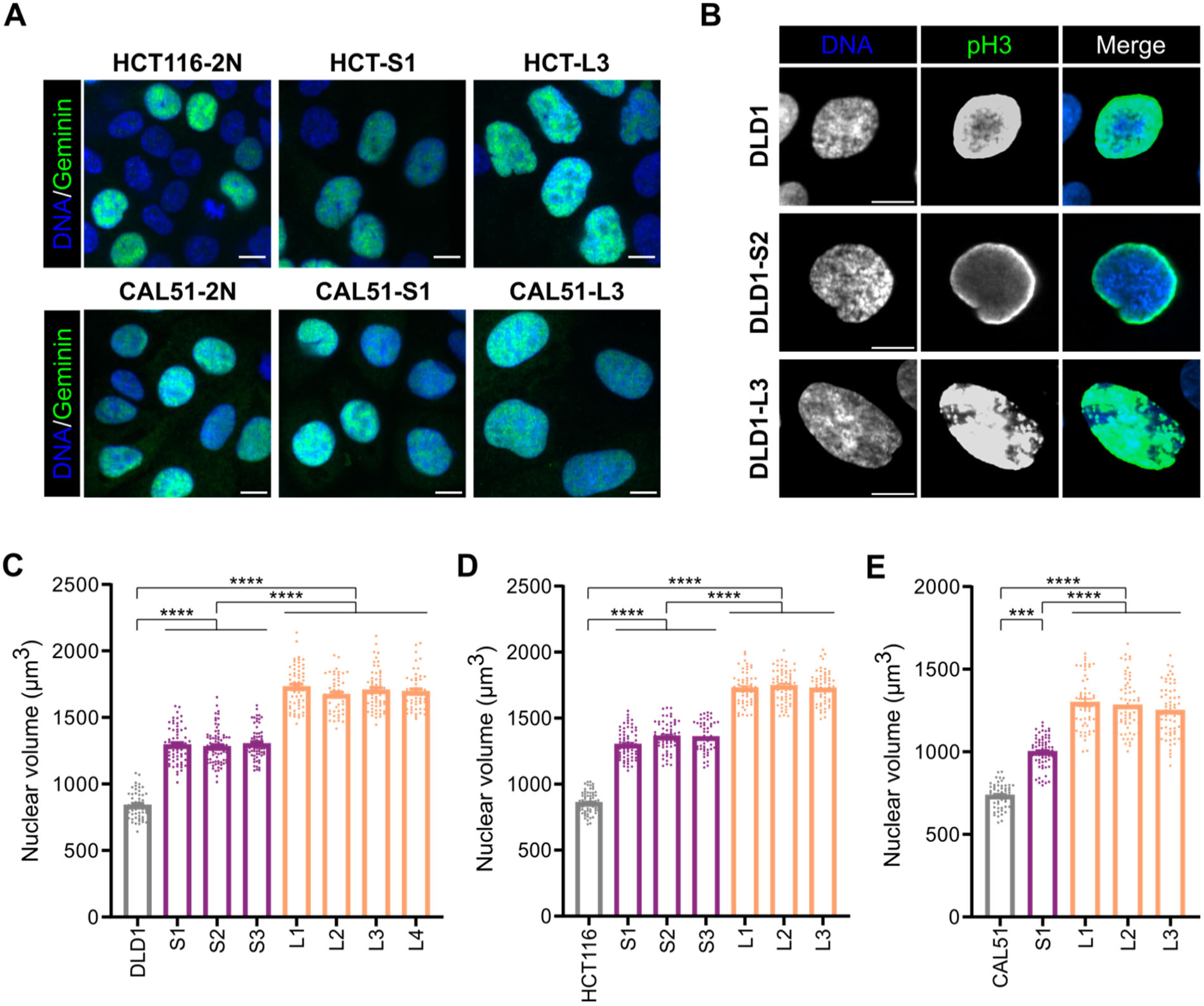
Nuclear volume differences in the S and L 4N clones are also observed at mitotic entry. **(A)** Representative images from HCT116 (top) and CAL51 (bottom) 2N cells (left) and S (middle) and L (right) 4N clones used for nuclear volume analysis in Figure 1D and E, respectively. **(B)** Representative images from 2N (top), S 4N (middle), and L 4N (bottom) DLD1 cells in prophase as determined by the presence of phosphorylated histone H3 (shown in green). **(C-E)** Quantification of nuclear volume in prophase cells from the DLD1, HCT116, and CAL51 2N cells, S 4N clones, and L 4N clones. Data are reported as mean ± SEM with individual points from two independent experiments (n ≥ 25 nuclei per experiment). All scale bars = 10 μm. *** = p<0.001, **** = p<0.0001.

**Supplementary Figure 2.**
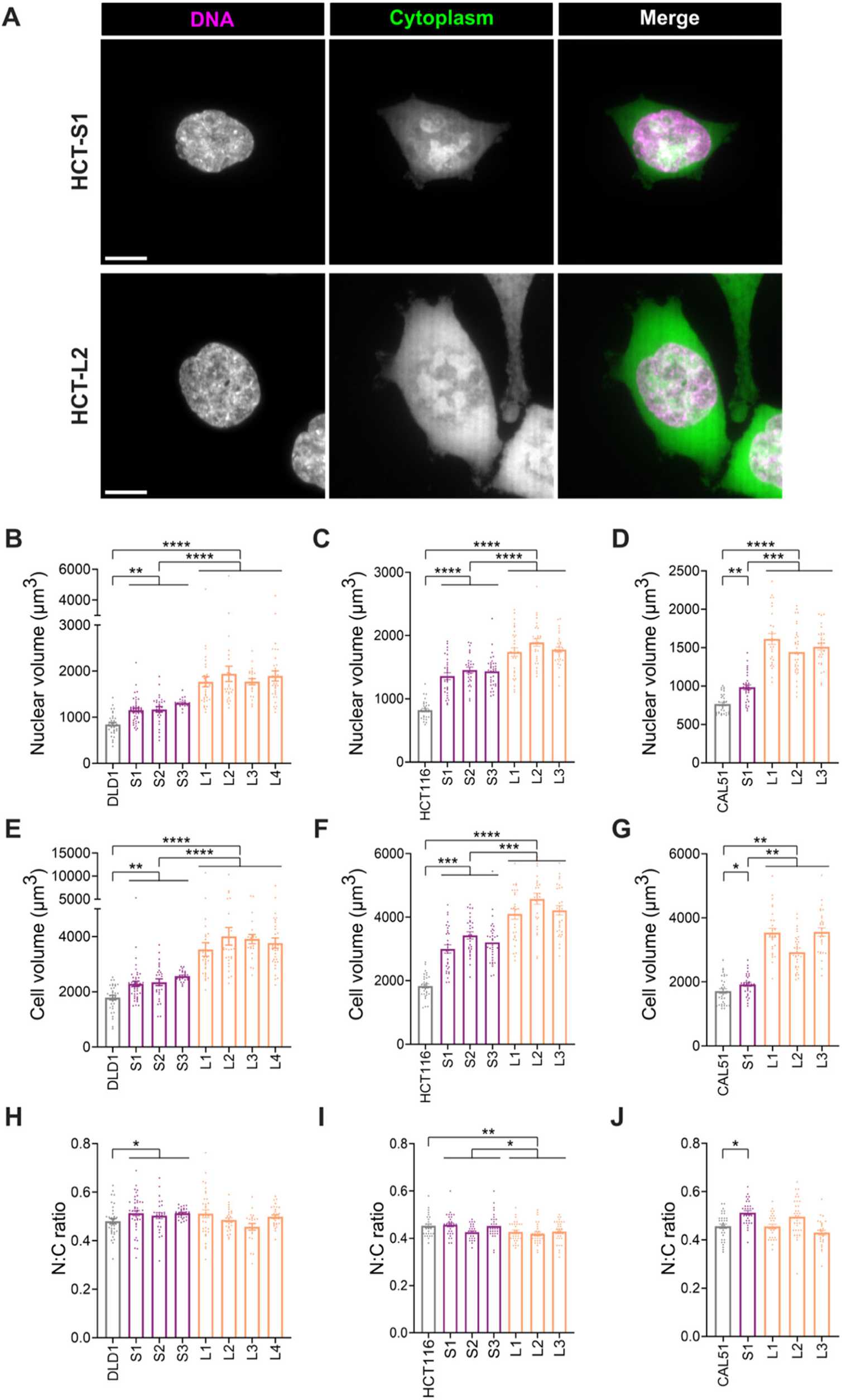
Cell volume, like nuclear volume, does not always scale with DNA content after WGD. **(A)** Representative images from HCT116 S1 (top) and L2 (bottom) used for paired nuclear (magenta) and cell (green) volume analysis within single cells. Quantification of **(B-D)** nuclear volume, **(E-G)** cell volume, and **(H-J)** N:C ratio in parental 2N cells and 4N clones from DLD-1, HCT-116, and CAL51 cell lines without the use of a cell cycle marker to exclude G1 cells. Data reported as mean ± SEM with individual points from at least two independent experiments (n ≥ 10 cells per experiment). All scale bars = 10 μm. * = p<0.05, ** = p<0.01, *** = p<0.001, **** = p<0.0001.

**Supplementary Figure 3.**
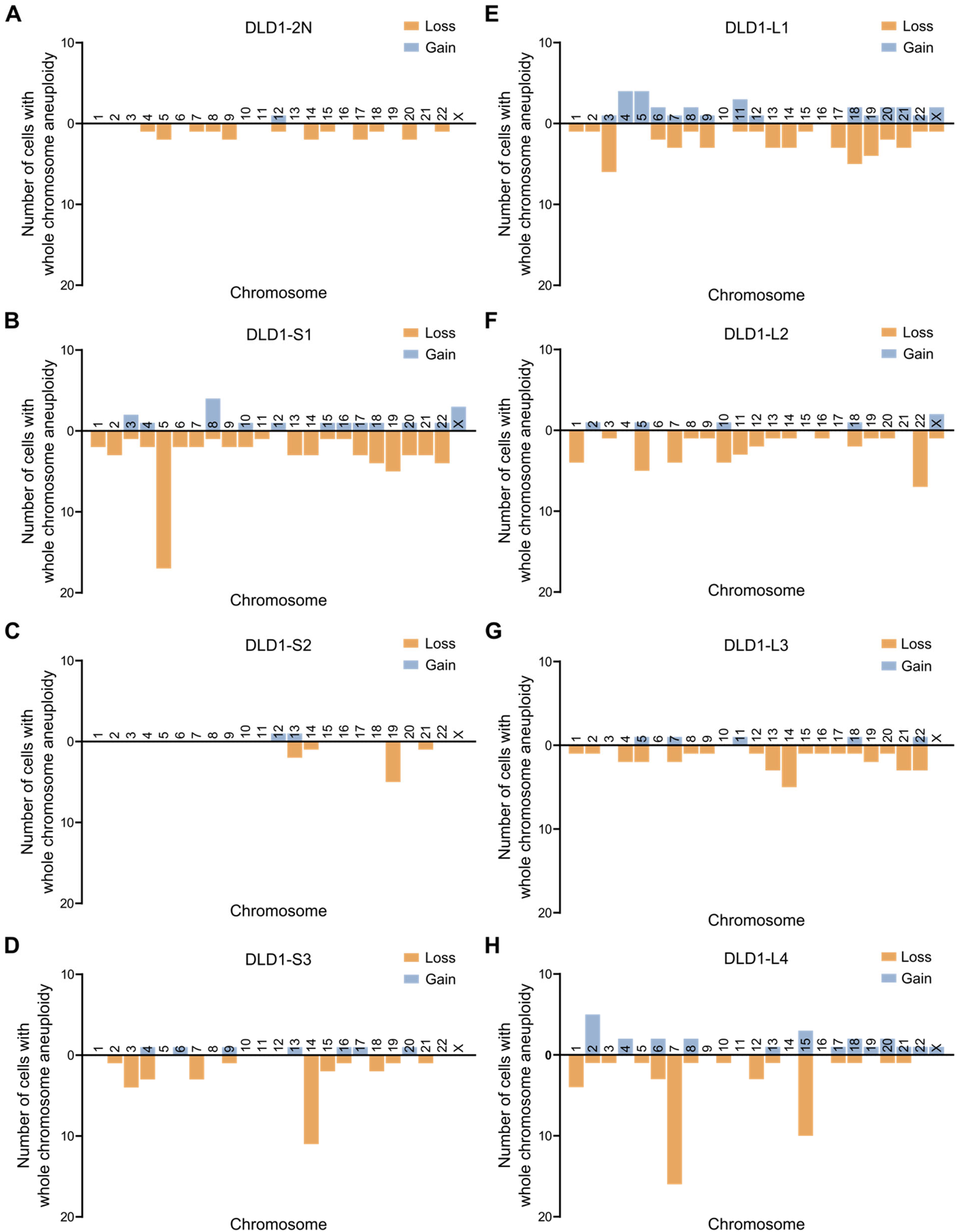
The S and L 4N DLD1 clones do not harbor recurrent chromosome copy number alterations. **(A-H)** Quantification of the number of cells with a whole chromosome gain or loss, based on mFISH, in the **(A)** 2N parental cells, **(B-D)** S 4N clones, and **(E-H)** L 4N clones (20 cells per clone).

**Supplementary Figure 4.**
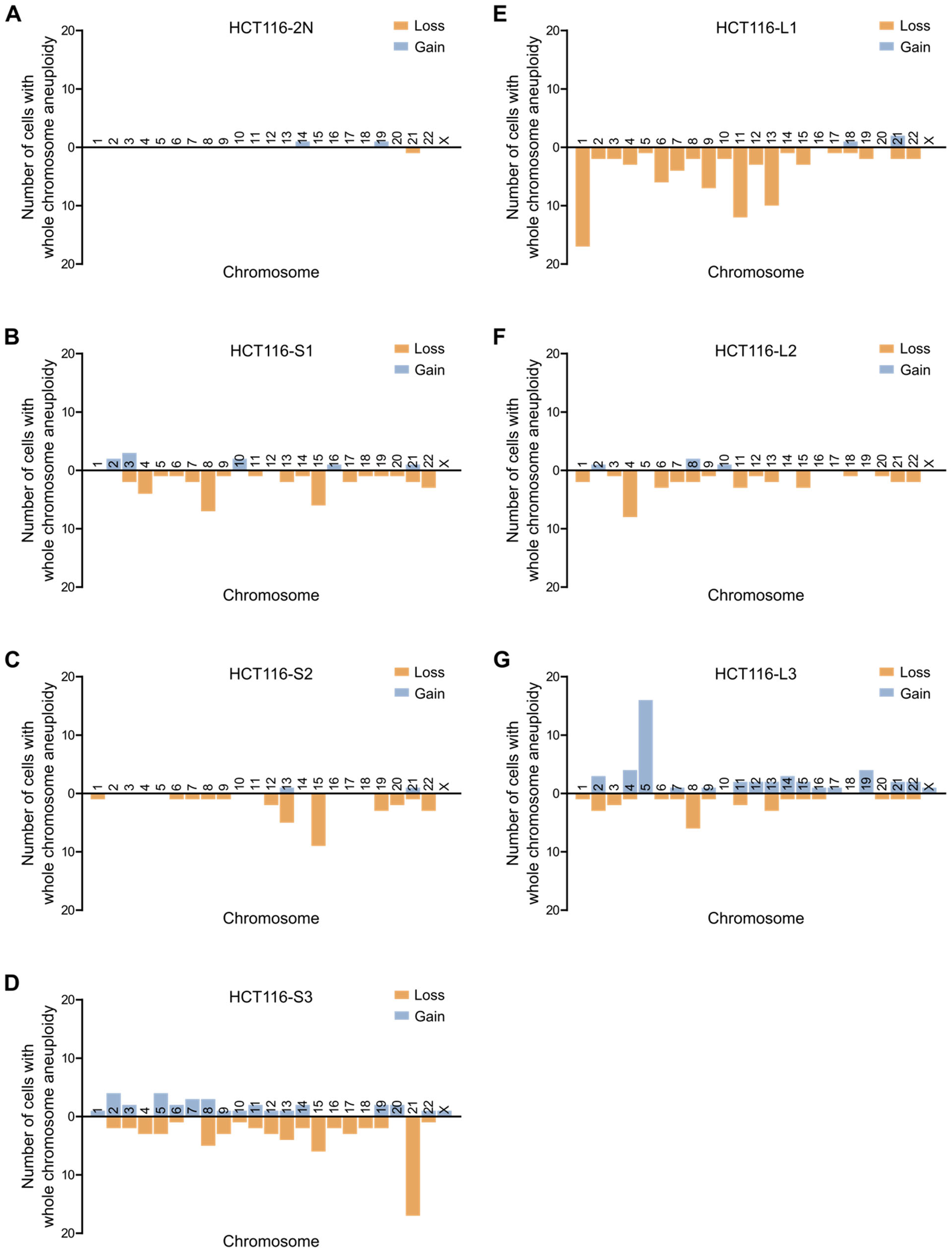
The S and L 4N HCT116 clones do not harbor recurrent chromosome copy number alterations. **(A-G)** Quantification of the number of cells with a whole chromosome gain or loss, based on mFISH, in the **(A)** 2N parental cells, **(B-D)** S 4N clones, and **(E-G)** L 4N clones (20 cells per clone).

**Supplementary Figure 5.**
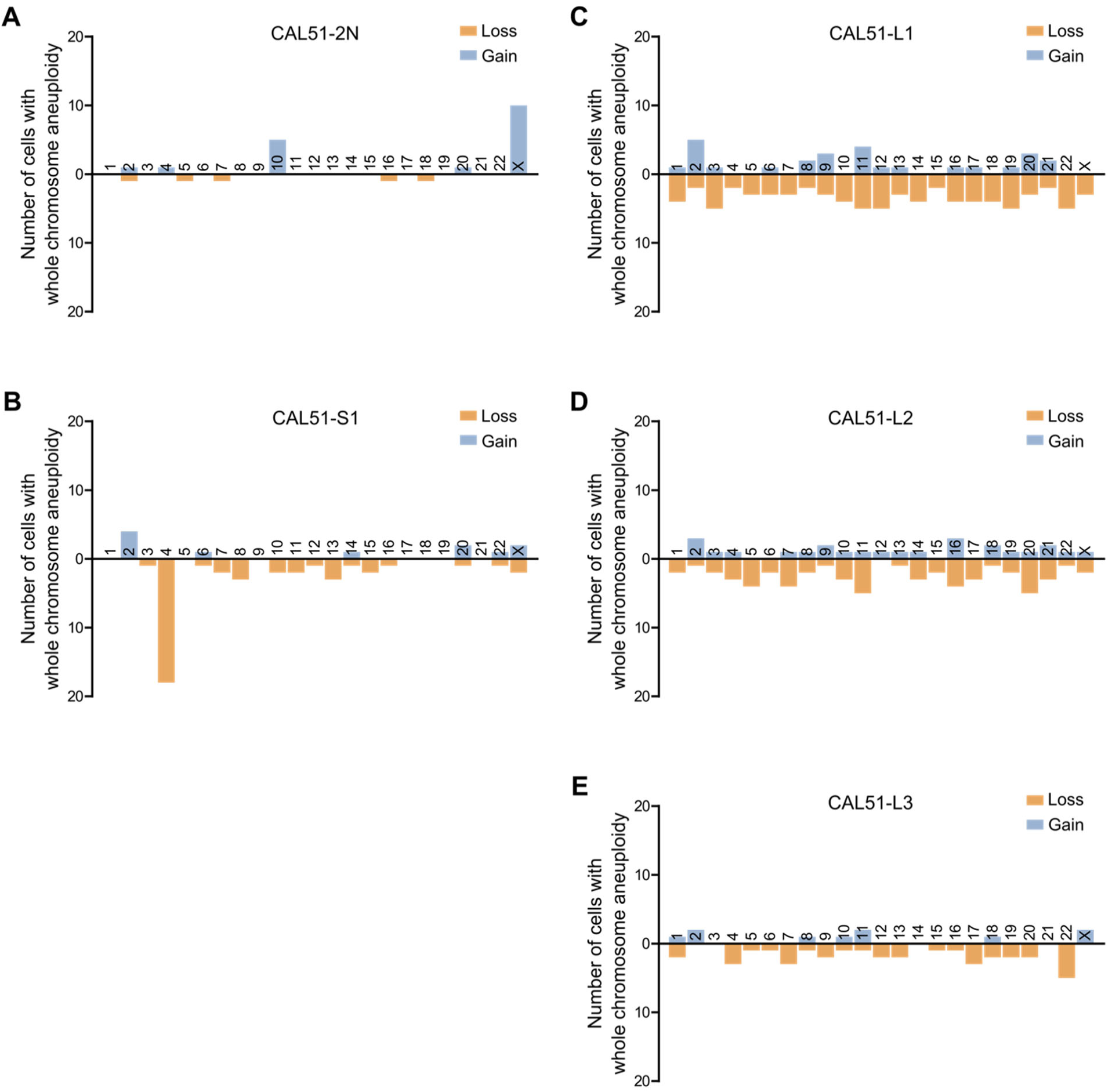
The L 4N CAL51 clones do not harbor recurrent chromosome copy number alterations. **(A-E)** Quantification of the number of cells with a whole chromosome gain or loss, based on mFISH, in the **(A)** 2N parental cells, **(B)** S 4N clone, and **(C-E)** L 4N clones (20 cells per clone).

**Supplementary Figure 6.**
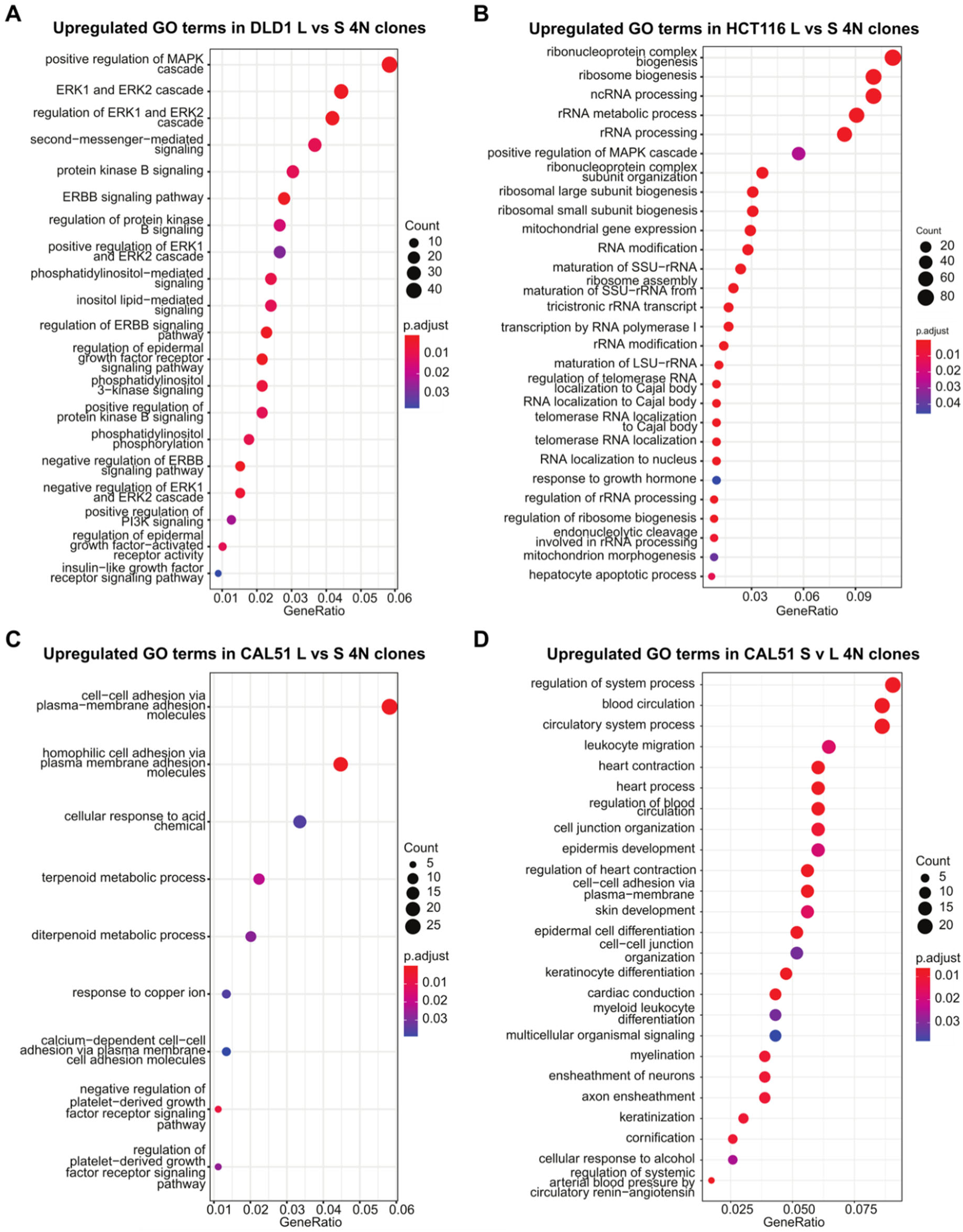
Pathways associated with cell growth, cell-cell adhesion, and biosynthesis are upregulated in the L compared to the S 4N clones. Dot plots showing the top GO terms enriched in the differentially expressed genes from **(A)** DLD1 L vs. S 4N, **(B)** HCT116 L vs. S 4N, **(C)** CAL51 L vs. S 4N, and **(D)** CAL51 S vs. L 4N. Corrected P-value of 0.05 and absolute fold change of 2 were set as the threshold for significant differential expression. GO terms with corrected p-value less than 0.05 were considered significant.

**Supplementary Figure 7.**
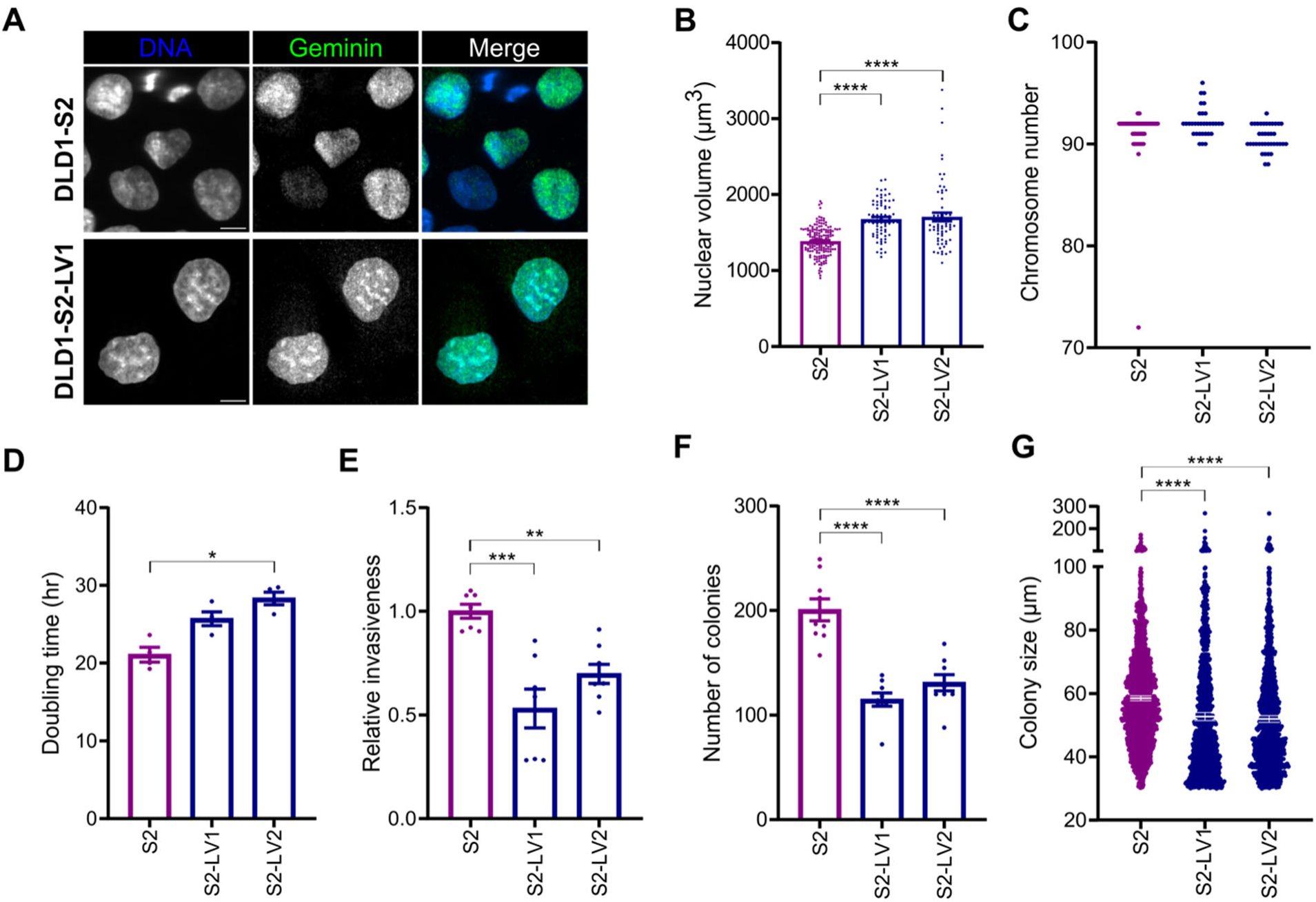
Increasing the nuclear size in a DLD1 S 4N clone affects cell phenotypes. **(A)** Representative images from DLD1-S2 (top) and DLD1-S2-LV1 (bottom) used for nuclear volume analysis. All scale bars = 10 μm. **(B)** Nuclear volume measurements in cells expressing geminin (green nuclei in panel A) from DLD1-S2 (magenta) and large variant clones LV1 and LV2 (blue). Data are reported as mean ± SEM with individual points from at least two independent experiments (n ≥ 30 nuclei per experiment). **(C)** Quantification of chromosome numbers in metaphase spreads from DLD1-S2, LV1, and LV2 cells (n ≥ 30). Quantification of **(D)** doubling times, **(E)** cellular invasion, and the **(F)** number and **(G)** size of colonies after 3 weeks in soft agar. Bar graphs report mean ± SEM for each group with individual data points showing the values of experimental replicates (n = 3) in **(D)** and technical replicates from **(E)** two and **(F)** three independent experiments. Individual data points in **(G)** show the major axis length from all soft agar colony measurements, and error bars depict the median ± 95%CI. Statistical significance was assessed using two-sample t-test to compare DLD1-S2 to its size variant clones. * = p<0.05, ** = p<0.01, *** = p<0.001, **** = p<0.0001.

**Supplementary Figure 8.**
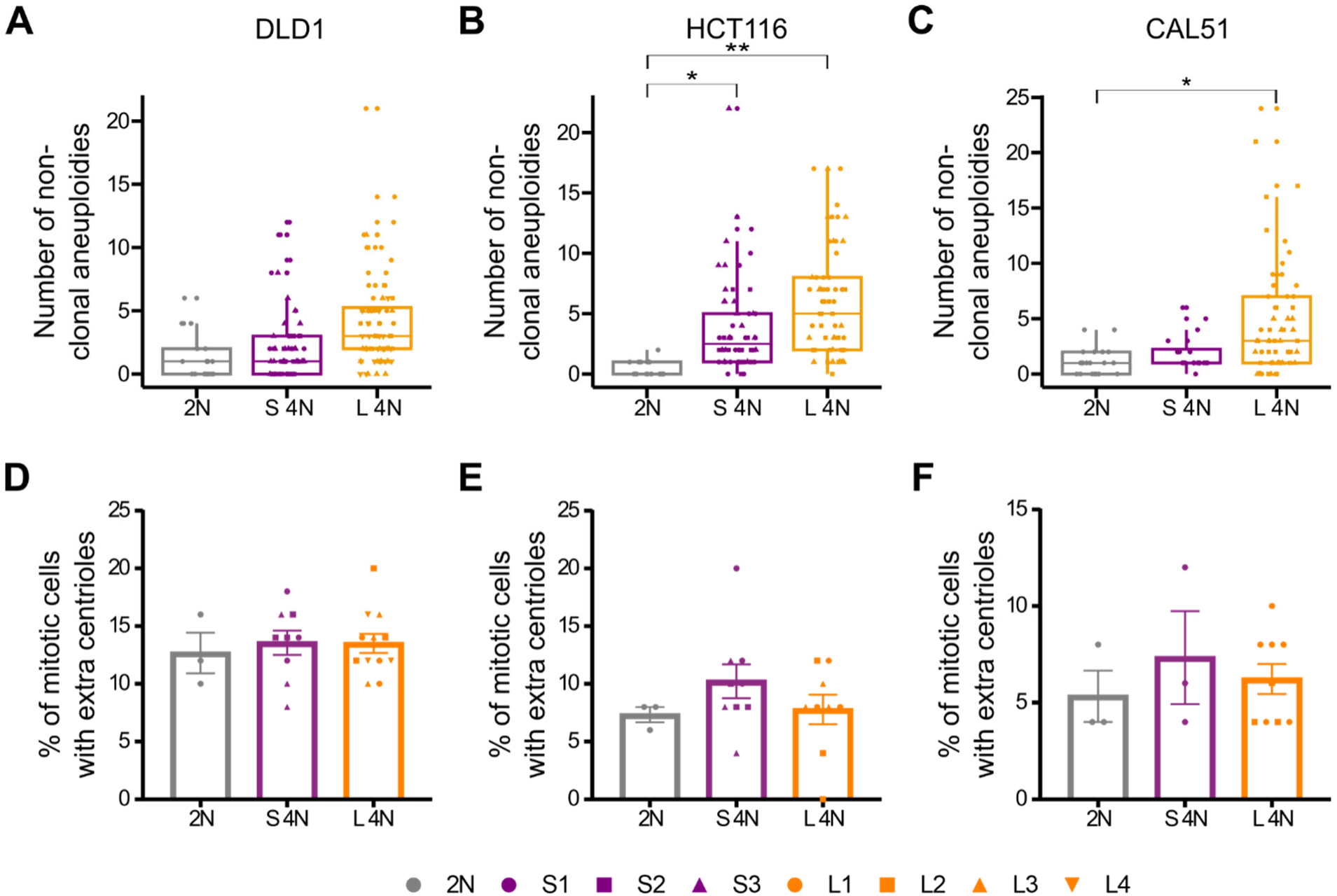
Chromosomal instability in the L 4N clones is not due to the presence of cells with extra centrioles. **(A-C)** Quantification of the total number of non-clonal whole chromosome and structural aneuploidies per karyotype in the 2N, S 4N, and L 4N clones. Each of the individual data points corresponds to a single karyotyped cell from each group. Box plots show the interquartile range (IQR) with a line at the median and whiskers extending to smallest and largest non-outlier values (defined as those within 1.5xIQR). **(D-F)** Quantification of centriole numbers in mitotic cells from the 2N, S 4N, and L 4N groups. Data depicted as mean ± SEM for each group with individual points showing the means of experimental replicates (n = 3) from each 2N cell line and 4N clone. 50 cells were analyzed per experiment.

**Supplementary Figure 9.**
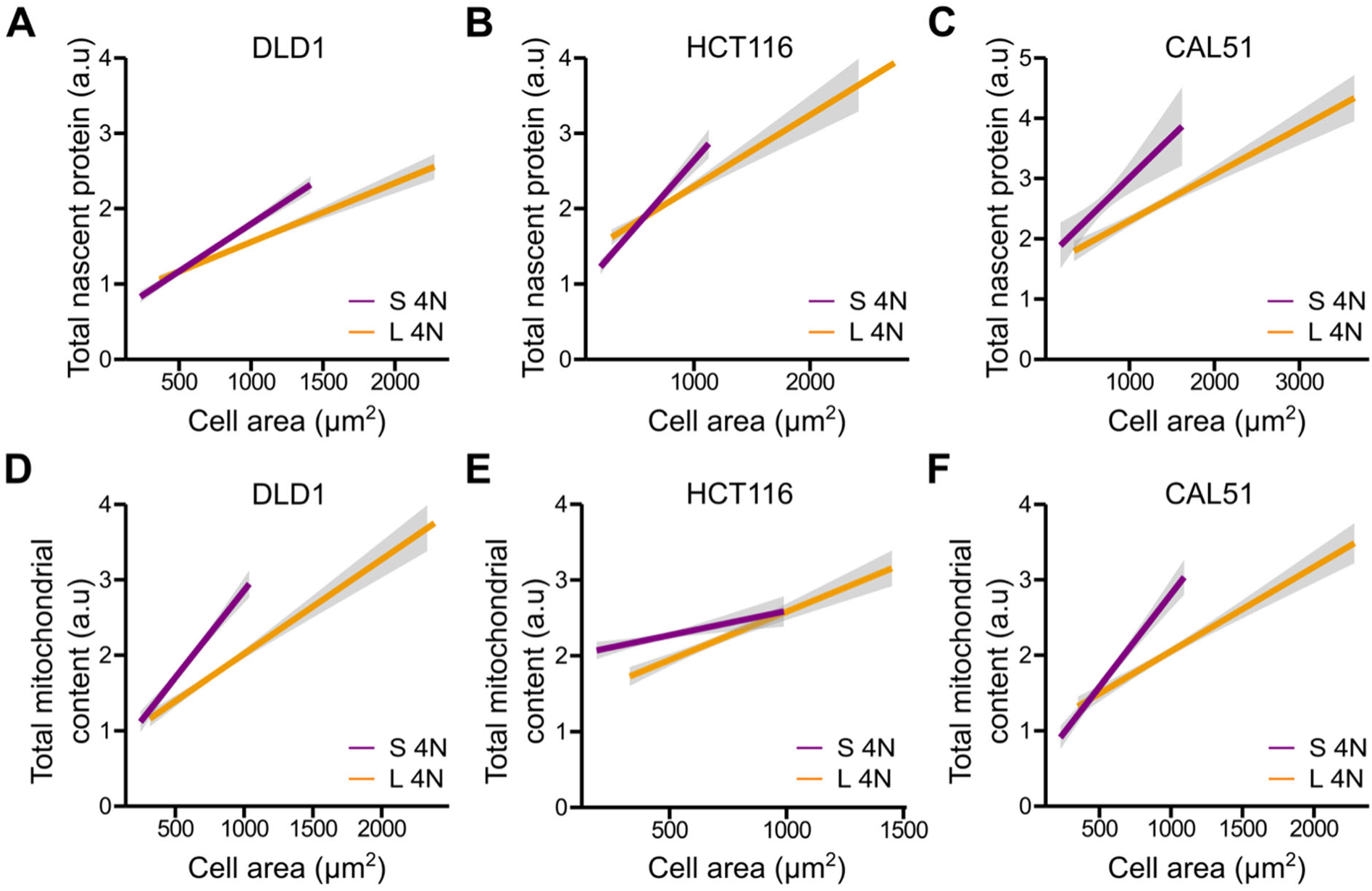
Protein synthesis and mitochondrial content tend to increase with cell size more rapidly in the S compared to the L 4N clones. Single cell OPP and MitoTracker intensities (from Figure 4) shown as a linear regression displaying the relationship between cell area and **(A-C)** total nascent protein levels as well as **(D-F)** total mitochondrial content in the S and L 4N clones from each cancer cell line.

**Supplementary Figure 10.**
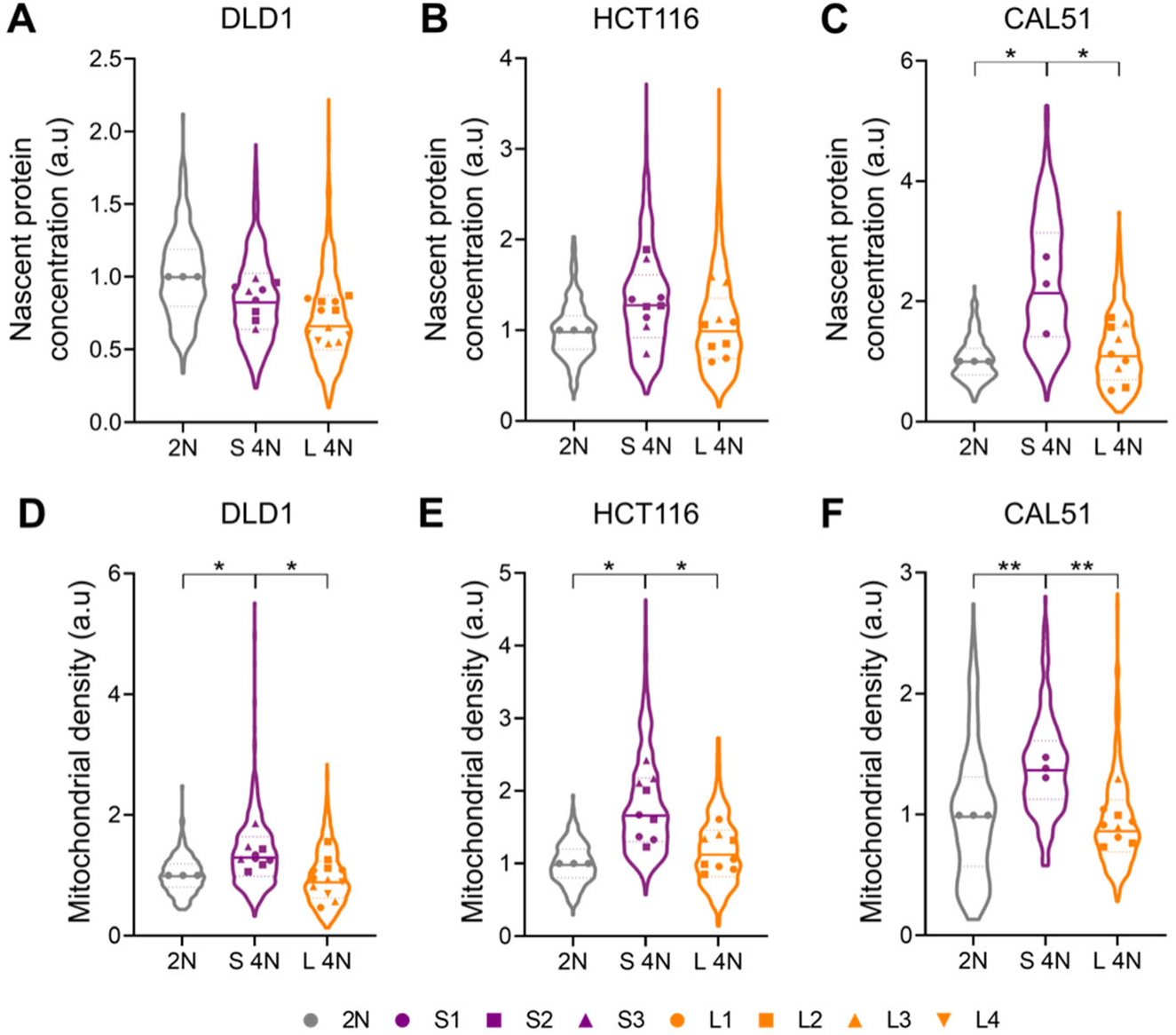
Nascent protein concentration and mitochondrial density per cell tend to be higher in the S than the L 4N clones. Quantification of **(A-C)** nascent protein concentration and **(D-F)** mitochondrial density (measured as mean fluorescence intensities). In all graphs, individual data points depict the means of experimental replicates (n = 3) for the 2N cells and 4N clones. Violin plots display the entire data distribution for the 2N, S 4N, and L 4N groups. Bold and dashed lines correspond to the median and quartiles, respectively. * = p<0.05, ** = p<0.01.

**Supplementary Figure 11.**
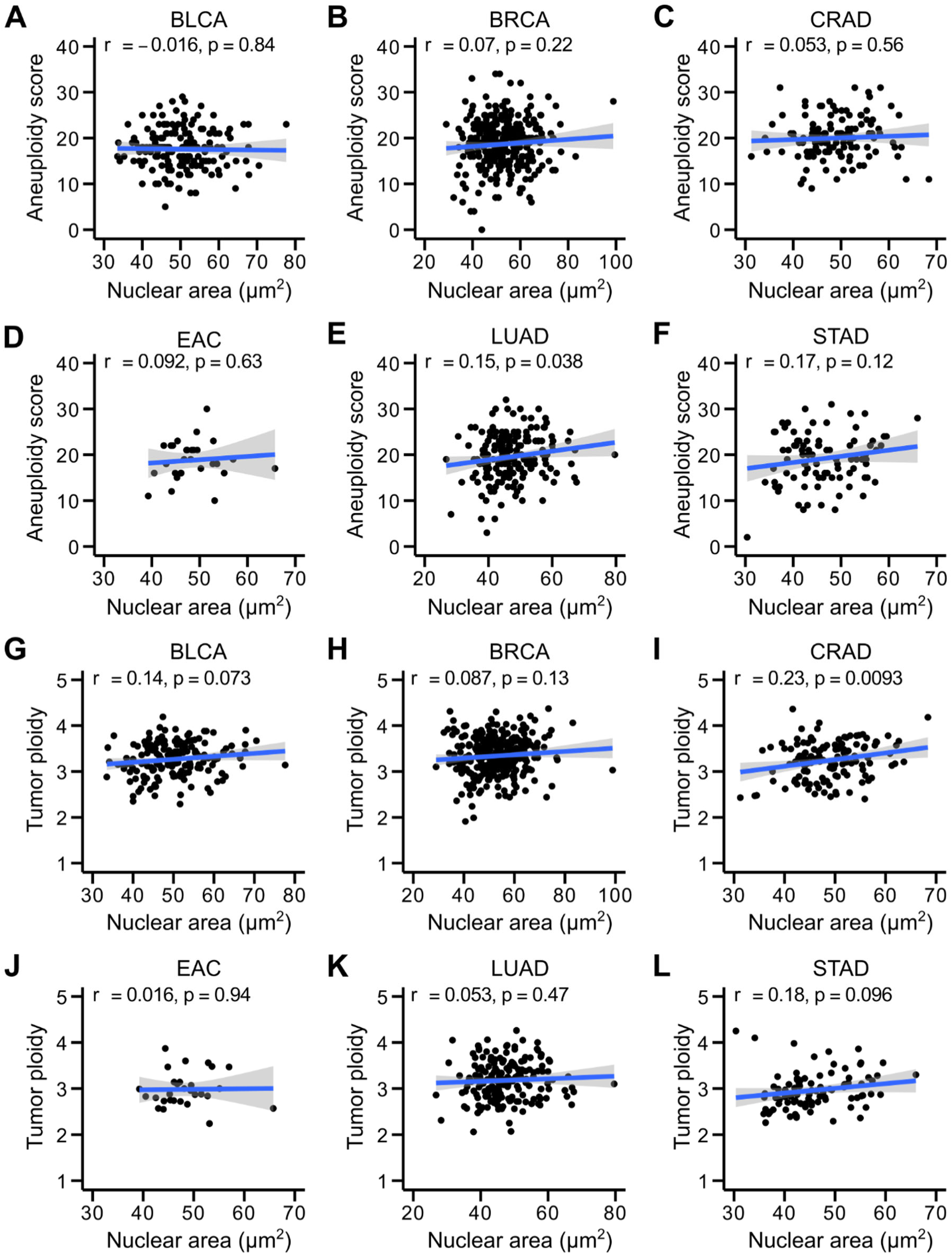
Neoplastic cell nuclear area does not correlate with aneuploidy score in WGD+ cancers in most tumor types. Pearson’s correlation analysis was used to assess the relationship between neoplastic cell nuclear size and **(A-F)** aneuploidy score and **(G-L)** tumor ploidy in WGD+ cancers from each tumor type.

**Supplementary Figure 12.**
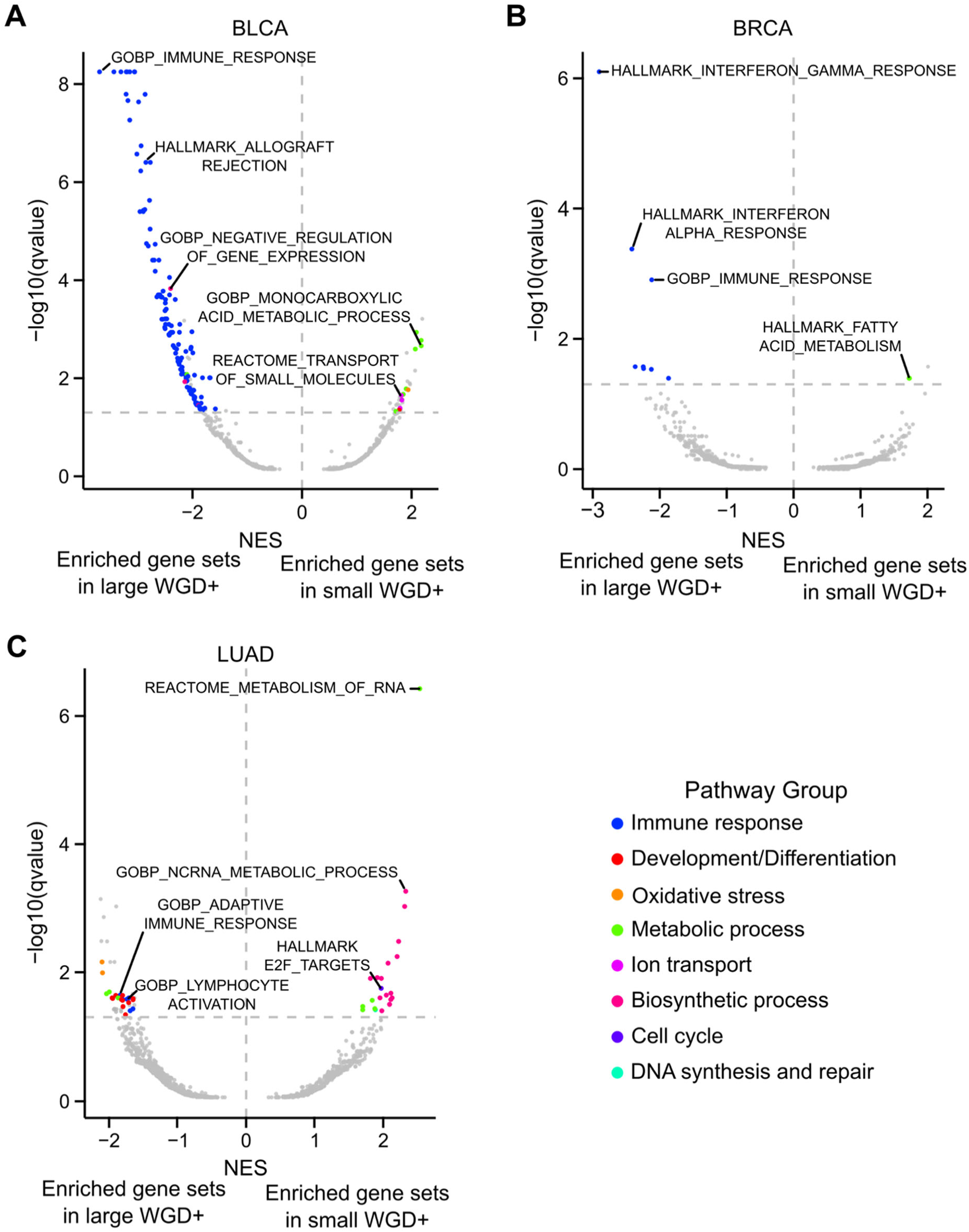
Cancer cell nuclear size is associated with distinct transcriptional signatures in WGD+ tumors. Comparison of the differential gene expression patterns (pre-ranked GSEA results) between the large and small WGD+ cancers in **(A)** BLCA, **(B)** BRCA, and **(C)** LUAD tumors. Individual points represent GO, KEGG, Reactome, and Hallmark gene sets. Significance threshold set at q-value=0.05 (horizontal lines). Recurrent pathway groups are color-coded.

